# Enhanced Production of Mesencephalic Dopaminergic Neurons from Lineage-Restricted Human Undifferentiated Stem Cells

**DOI:** 10.1101/2021.09.28.462222

**Authors:** Muyesier Maimaitili, Muwan Chen, Fabia Febbraro, Noëmie Mermet-Joret, Johanne Lauritsen, Ekin Ucuncu, Ida Hyllen Klæstrup, Per Qvist, Sadegh Nabavi, Marina Romero-Ramos, Mark Denham

## Abstract

The differentiation of human pluripotent stem cells (hPSCs) into mesencephalic dopaminergic (mesDA) neurons requires a precise combination of extrinsic factors that recapitulates the *in vivo* environment and timing. Current methods are capable of generating authentic mesDA neurons after long-term culture *in vitro*; however, when mesDA progenitors are transplanted *in vivo*, the resulting mesDA neurons are only minor components of the graft. This low yield hampers the broad use of these cells in the clinic. In this study, we genetically modified pluripotent stem cells to generate a novel type of stem cells called lineage-restricted undifferentiated stem cells (LR-USCs), which robustly generate mesDA neurons. LR-USCs are prevented from differentiating into a broad range of nondopaminergic cell types by knocking out genes that are critical for the specification of cells of alternate lineages. Specifically, we target transcription factors involved in the production of spinal cord and posterior hindbrain cell types. When LR-USCs are differentiated under caudalizing condition, which normally give rise to hindbrain cell types, a large proportion adopt a midbrain identity and develop into authentic mesDA neurons. We show that the mesDA neurons are electrophysiologically active, and due to their higher purity, are capable of restoring motor behavior eight weeks after transplantation into 6-hydroxydopamine (6-OHDA)-lesioned rats. This novel strategy improves the reliability and scalability of mesDA neuron generation for clinical use.

## Introduction

Mesencephalic dopaminergic (mesDA) neurons develop from the ventral midbrain of the neural tube. The morphogen sonic hedgehog (SHH) and members of the WNT family are instrumental in their specification due to their essential role in establishing the dorsal-ventral and anterior-posterior (A-P) axes of the embryo, respectively ^1,2^. For the production of midbrain dopaminergic (DA) neurons, high SHH signaling from the notochord is required to specify ventral neural epithelial cells in the neural plate to a floor plate identity ^3^, and graded Wnt signaling emanating from the posterior regions of the embryo is required to specify anterior neuroectoderm to a midbrain identity ^4^. At later stages in development, neural progenitors in the midbrain region receive Wnt1 and Fgf8 signals from the isthmic organizer, which refines the patterning of the cells to a caudal location within the mesencephalon ^5,6^. Recapitulating these developmental steps *in vitro* with human pluripotent stem cells (hPSCs) is the goal of stem cell transplantation therapies for Parkinson’s disease ^7^.

In stem cell differentiation protocols, early application of high concentrations of SHH is necessary to specify neural progenitor cells to a floor plate identity ^8,9^. However, along the A-P axis (also referred to as rostral-caudal axis), specification to a caudal midbrain identity is more complex. A titrated WNT concentration within a precise range can specify anterior neuroectoderm progenitors to a caudal midbrain identity ^10,11^. High concentrations result in the production of hindbrain cell types, and lower concentrations result in an anterior midbrain or diencephalic identity. Timed delivery of FGF8 or sequential exposure to high levels of WNT has also been shown to improve specification to a caudal midbrain identity ^12,13^. Despite these advances in stem cell differentiation protocols, the overall yield of mesDA neurons after transplantation is extremely low, and increasing the yield would significantly improve cell purity and the reliability of graft function, which is essential for the use of these cells in the clinic ^14,15^.

Downstream of extrinsic signaling, progenitor cells employ gene regulatory networks (GRNs) to interpret morphogen gradients and regulate cell fate ^16^. During neural tube development, Otx2 is expressed in anterior regions, whereas Gbx2 is expressed in posterior regions at early stages of development. The midbrain-hindbrain boundary demarcates the Otx2 and Gbx2 boundary and the location of the isthmic organizer ^6^. At this border, Otx2 and Gbx2 establish a separate network of transcription factors that maintain the position of the isthmic organizer and assist in patterning the surrounding region ^5,17,18^. Alterations in the expression levels of transcription factors in networks can result in the expansion or loss of specific brain regions ^19^.

Transcription factor networks have been altered by forced expression of lineage-determinant transcription factors, such as Lmx1a, to accelerate the differentiation of mouse embryonic stem cells (ESCs) and human embryonic stem cells (hESCs) into DA cells ^20,21^, and forced expression of *GLI1* in human ESC-derived neural progenitors can generate floor plate cells ^22^. Conversely, in developing embryos, ablation of transcription factors can result in the loss of specific cell populations. Along the A-P axis, deletion of *Otx2* in embryos results in the loss of forebrain and midbrain structures ^23,24^. Null mutations in all three *Cdx* family members result in the loss of spinal cord cell types below the preoccipital level due to disruption of central and posterior *Hox* gene expression and prevention of neuromesodermal progenitor (NMP) formation ^25–27^. Interestingly, null mutations in *Gbx2* result in expansion of the midbrain at the expense of rhombomeres (r)1-3 ^28^. These studies demonstrate that ablation of transcription factors that control cell fate can lead to the activation of altered GRNs and the respecification or expansion of alternate populations. It is therefore possible that transcription factor determinants in GRNs that are involved in lineage choices can be disrupted to control cell fate and bias the differentiation of hPSCs toward a mesencephalic neuron identity.

In this study, we used a gene knockout approach to restrict cell fate and prevent the differentiation of non-DA cell lineages with the aim of enhancing differentiation to mesDA neurons. Specifically, we focused on the early developmental stages when major lineage choices are made and identified the transcription factor determinates that are critical for those lineages but not required for a mesDA fate. By inducing loss-of-function mutations in lineage determinant genes expressed in non-DA lineages, we were able to bias the differentiation of hPSCs toward a mesDA identity. We generated stem cells that could be expanded in the undifferentiated pluripotent state and were restricted in their potential when differentiated. We named these lineage-restricted undifferentiated stem cells (LR-USCs). Specifically, we focused on the A-P axis of the developing neural tube because mesDA progenitors require titrated expression of WNT and because regulating this axis would most benefit a mesDA neuron differentiation protocol. We examined transcription factors that regulate hindbrain and spinal cord cell fates. The anterior hindbrain has been shown to require Gbx2, and the spinal cord is dependent on Cdx family members ^26,28^. By ablating these genes, we increased the percentage of cells that adopted a midbrain identity and prevented specification to a spinal cord fate. Uniquely, LR-USCs generated mesDA neurons under unfavorable conditions, specifically high concentrations of WNT, which normally induce the generation of hindbrain and spinal cord cell types. Furthermore, we demonstrated that the mesDA neurons were functional and able to restore motor behavior in a rodent model of 6-hydroxydopamine (6-OHDA)-induced Parkinson’s disease. Our results show that this approach can be successfully used to robustly generate functional mesDA neurons and that the development of these cells is less reliant on specific concentrations of extrinsic factors. This strategy significantly improves the reliability and scalability of mesDA neuron production for clinical use for cell transplantation.

## Results

### Midbrain cell types are preferentially generated from PSCs containing homozygous null mutations in *GBX2, CDX1, CDX2, and CDX4* under caudalizing conditions

To restrict the differentiation of hPSCs and guide them toward a mesDA neuron identity, we introduced null mutations in transcription factors that regulate cell fate along the A-P axis. First, we investigated whether a homozygous null mutation in *GBX2* in hESCs (H9 cells) results in an increase in the production of midbrain cell types when the cells are differentiated under conditions known to produce hindbrain and spinal cord cells. We generated a *GBX2*^-/-^ hESC line by introducing indels into the coding sequence (Supplementary Fig. 1). In the undifferentiated state, the *GBX2*^-/-^ cell line was morphologically indistinguishable from control hESCs and capable of differentiating into neural progenitors. Using our previously published protocol for generating caudal neural progenitors (CNPs) ^29^, we differentiated *GBX2*^-/-^ cells into CNPs for four days *in vitro* (DIV) and compared these cells to H9 (control) CNPs (Fig. 1a-b). Indeed, when differentiated in the presence of a GSK3B inhibitor (GSK3i; CHIR99021) at a concentration known to give rise to hindbrain and spinal cord cells (3 µM), there was a small but significant increase in the expression of the forebrain/midbrain marker *OTX2* (LogFC to hESC = H9: 0.003; *GBX2*^-/-^: 0.083; P = 0.0005) (Fig. 1c) and a significant reduction in the transcript level of *CDX2* in the *GBX2*^-/-^ cells compared to the H9 cells (LogFC to hESC = H9: 8871.79; *GBX2*^-/-^: 3160.57; P = 0.006) (Fig. 1d). Despite the increase in *OTX2* expression, we observed few OTX2-positive cells, and almost all cells were CDX2-positive (Fig. 1b).

**Figure 1:**
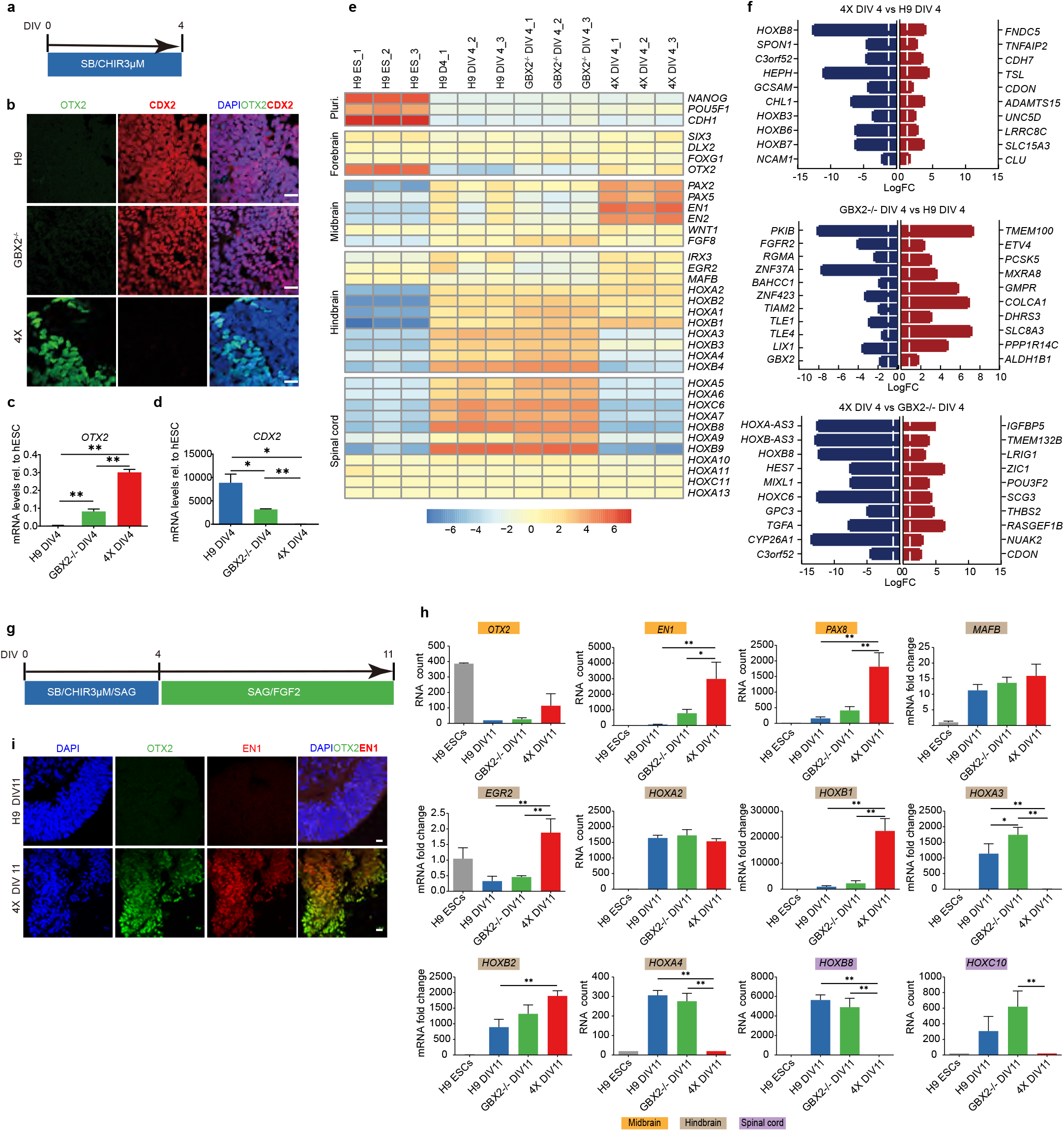
Differentiation of GBX2^-/-^ and 4X cells into CNPs. **a**, Schematic diagram of the 4 DIV CNP differentiation protocol. **b**, Representative immunostaining images of H9 and GBX2^-/-^ CNPs at 4 DIV showing OTX2^-^/CDX2^+^ cells. A few 4X CNPs were positive for OTX2, but no cells were positive for CDX2. Scale bars, 20 µm. **c-d**, qPCR analysis of *OTX2* (c) and *CDX2* (d) expression in H9, GBX2^-/-^ and 4X CNPs at 4 DIV. The data are presented as the mean ± SD; n= 3. One-way ANOVA showed statistical significance, and then an unpaired t-test comparing two groups was performed. *P< 0.05; **P< 0.01. **e**, Heatmap of the expression of pluripotent and neural genes representing the anterior, midbrain, hindbrain and spinal cord regions in H9, GBX2^-/-^ and 4X CNPs at 4 DIV. **f**, The top 10 downregulated (blue) and upregulated (red) genes (and additional selected genes in bold) between cultured 4X and H9 cells, cultured GBX2^-/-^ and H9 cells and cultured 4X and GBX2^-/-^ cells at 4 DIV. The threshold bar (white line) indicates a fold change of ±2. **g**, Schematic diagram of the 11 DIV CNP differentiation protocol. **h**, RNA expression analysis of the midbrain genes (orange) *OTX2, EN1* and *PAX8*; the hindbrain genes (gray) *MAFB, EGR2, HOXA2, HOXB1, HOXA3, HOXB2*, and *HOXA4*; and the spinal cord genes (purple) *HOXB8* and *HOXC10*. The data are presented as the mean ± SD; n= 3. One-way ANOVA followed by Tukey’s multiple comparisons test. *P< 0.05; **P< 0.01. **i**, Representative immunohistochemical analysis of OTX2/EN1 double-positive cells among 11 DIV 4X cells. No OTX2/EN1 double-positive cells were detected among H9 cells. Scale bars, 10 µm. For (b) and (i), DAPI was used as a nuclear stain. Caudal neural progenitor: CNP.

Based on the results obtained using the *GBX2*^-/-^ line, we next attempted to further restrict the potential of cells along the A-P axis by knocking out *CDX* family members. Cdx2 is an upstream regulator of central and posterior *Hox* genes and acts as a key determinant of spinal cord fate through its regulation of axial elongation ^25,30^. Triple homozygous null mutations in all three *Cdx* genes (*Cdx1, Cdx2*, and *Cdx4*) result in severe truncation of the spinal cord below the postoccipital level and prevent the formation of NMPs ^26,27^. Because loss of Cdx1/2/4 in mice causes posterior truncation, we chose to disrupt the *CDX* family of genes in addition to knocking out *GBX2*. We generated homozygous null mutations in all three *CDX* family members, *CDX1/2/4*, by targeting their DNA binding domains (Supplementary Fig. 2). The resulting *GBX2*^*-/-*^*CDX1/2/4*^*-/-*^ hESC line (hereafter referred to as 4X cells) was differentiated for four DIV using the CNP protocol (Fig. 1a). As expected, we did not detect *CDX2* transcripts or CDX2 expression in 4X CNPs (Fig. 1b, d). Strikingly, at 4 DIV, the 4X neural progenitors showed a significant increase in *OTX2* transcript levels compared to H9 and *GBX2*^-/-^ derived CNPs (LogFC to hESC = H9: 0.0032; *GBX2*^-/-^: 0.083; 4X: 0.301; P < 0.0001 for both 4X vs. H9 or GBX2^-/-^), and we could readily identify OTX2-positive cells (Fig. 1b, c).

To elucidate the effects on gene expression in more detail, we performed RNA sequencing on CNPs of all three cell lines (H9, *GBX2*^-/-^ and 4X). We assessed *HOX* gene expression profiles and found that the 4X CNPs showed a significant reduction in the expression of posterior *HOX* genes, compared to H9, beginning with *HOXA3* and moving caudally (*HOXA3*, P = 1×10^−6^; *HOXA5*, P = 1.95×10^−5^; HOXA7, P = 0.0005; *HOXA9*, P = 0.0004; *HOXA10* P = 0.002; Fig. 1e). These results indicate that 4X cells were unable to generate progenitor cell types caudal to r4 ^31^. We next questioned whether there were changes in the expression of anterior genes. First, we examined the expression of forebrain genes in 4X CNPs and observed no change in the expression of *SIX3, DLX2* and *FOXG1* (Fig. 1e). However, the transcript levels of the forebrain/midbrain gene *OTX2* were significantly increased in 4X CNPs compared to H9 and GBX2^-/-^ CNPs (H9, LogFC = 6.83, P = 0.001; *GBX2*^-/-^, LogFC = 3.92, P = 0.003, respectively). The expression of the midbrain genes *PAX2*, and *EN1* was also significantly increased in 4X CNPs compared to H9 (*PAX2*, LogFC = 2.63, P = 0.03; *EN1*, LogFC = 4.76, P = 0.03) and *GBX2*^-/-^ CNPs (*PAX2*, LogFC = 3.49, P = 0.00005; *EN1*, LogFC = 7.08, P = 0.0003; Fig. 1e). Interestingly, *GBX2*^-/-^ cells showed a reduction in the expression of anterior hindbrain genes, such as *EGR2* (LogFC = -3.86; also known as *KROX20)* and *MAFB* (LogFC = -0.67), which was in line with reports showing that disruption of *Gbx2* in mice causes loss of r1-3 ^28^. In contrast, the expression of *MAFB* significantly increased in 4X cells (*MAFB*, LogFC = 2.49, P = 0.0002), which is in accordance with the loss of posterior *HOX* expression.

Analysis of differentially expressed genes among the three groups showed that the top significantly downregulated genes in 4X cells compared to H9 and *GBX2*^-/-^ cells included *HOX* genes (Fig. 1f). Furthermore, the expression of *CYP26A1*, which is involved in retinoic acid (RA) metabolism and is induced by CDX2, was significantly downregulated in 4X cells compared to *GBX2*^-/-^ cells (LogFC = -13.43, P = 1.30×10^−12^). A comparison of *GBX2*^-/-^ to H9 cells showed that knockout of GBX2 alone resulted in a significant decrease in the transcription levels of the Groucho corepressor proteins TLE1 (LogFC = -2.86, P = 8.05×10^−9^) and TLE4 (LogFC = -1.54, P = 8.32×10^−9^), which function with GBX2 to repress OTX2 ^32^. Overall, knockout of *GBX2* resulted in disruption of anterior hindbrain patterning, and 4X cells showed that further loss of CDX family members caused a posterior limitation of the CNS equivalent of r4, which resulted in significantly higher expression of midbrain and anterior hindbrain genes (Fig. 1e, f).

To further explore the potential of the 4X cells, we extended the duration of differentiation to 11 DIV and added smoothened agonist (SAG) to ventralize the cells (Fig. 1g). We found that the transcript levels of *OTX2* were higher in the 4X cells than in the H9 and *GBX2*^-/-^ cells, which was similar to what was observed after differentiation for 4 DIV, although at 11 DIV it was not significant (RNA count, H9: 20.0; *GBX2*^-/-^: 26.9; 4X: 114.3; P = 0.09 and P = 0.12, respectively) (Fig. 1h). Furthermore, we found that the expression of the midbrain gene *PAX8* was significantly upregulated in 4X cells compared to H9 and *GBX2*^-/-^ cells (RNA count, H9: 158.6; *GBX2*^-/-^: 414.4; 4X: 1813.0; P = 0.0007, P = 0.002, respectively) (Fig. 1h). The expression of *EN1*, which spans the caudal midbrain and r1 during development ^33^, was also significantly upregulated in 4X cells compared to H9 and *GBX2*^-/-^ cells (RNA count = H9: 66.6; *GBX2*^-/-^: 784.8; 4X: 2985.4; P = 0.003 and P = 0.01, respectively; Fig. 1h). The expression of hindbrain gene *EGR2*, which is expressed in r3 and r5 ^34^, was significantly upregulated (LogFC = H9: 0.33; *GBX2*^-/-^: 0.46; 4X: 1.88; P = 0.001 and P = 0.002, respectively; Fig. 1h), and the expression of *MAFB*, a marker of r5 and r6 ^34^, was not significantly altered.

Upon examination of *HOX* expression profiles, we found that central and posterior *HOX* genes beginning with *HOXA3* and moving posteriorly were absent in 4X cells (Fig. 1h). The expression of *HOXA2*, which is expressed throughout the hindbrain (except for r1), was maintained in the 11 DIV 4X CNPs; however, the expression of the anterior *HOX* genes *HOXB2* and *HOXB1* was significantly upregulated in 4X cells compared to H9 cells (LogFC = H9: 893.4; 4X: 1893.3; P = 0.005 for *HOXB2*, and H9: 965.0; 4X: 22377.6; P = 0.0002 for *HOXB1*), suggesting a compensatory shift in the population to a more anterior identity (Fig. 1h). Immunocytochemical analysis confirmed the change that we observed at the transcript level (Fig. 1i). We identified caudal midbrain progenitors, i.e., OTX2/EN1 double-positive cells, among 4X cells at 11 DIV, but such cells were not detected among H9 cells at 11 DIV (Fig. 1i). These results indicate that under caudalizing conditions, 4X cell did not produce spinal cord progenitors and showed a restricted *HOX* expression profile up to r4. Furthermore, the distribution of cell types was shifted anteriorly, as indicated by the presence of OTX2/EN1 double-positive cells, which were not detected among H9 cells. These results support the notion that 4X cells preferentially adopt a midbrain or anterior hindbrain identity under conditions that usually give rise to caudal hindbrain and spinal cord cell types.

### LR-USCs efficiently generate caudal midbrain progenitors

We next investigated how 4X cells respond when differentiated with a range of GSK3i concentrations and whether the enhanced specification of 4X cells to OTX2/EN1 double-positive midbrain cells at 3 µM GSK3i is maintained at lower GSK3i concentrations. Using the same 11 DIV caudal protocol, we tested four concentrations of GSK3i from 0.7 µM to 3 µM (Fig. 2a-b). We examined the percentage of cells expressing OTX2 by flow cytometry. Strikingly, at the lowest concentrations of GSK3i, i.e., 0.7 µM and 1 µM, 71.0% and 34.9%, respectively, of 4X cells were OTX2-positive; there were significantly more OTX2-positive 4X cells than OTX2-positive H9 and GBX2^-/-^ cells at these concentrations (H9: 2.5%; GBX2^-/-^: 21.8%; 4X: 71.0% P < 0.0001 for both, for 0.7 µM. H9: 4.8%; GBX2^-/-^: 5.0%; 4X: 34.9% P < 0.0001 for both, for 1 µM). At 2 - 3 µM GSK3i, the percentage of OTX2-positive cells decreased dramatically in all groups; however, there were still more OTX2-positive 4X cells than OTX2-positive H9 and GBX2^-/-^ cells at 3 µM GSK3i (H9: 0.3%; GBX2^-/-^: 0.8%; 4X: 10.1% P < 0.0004 and P < 0.0008, respectively).

**Figure 2:**
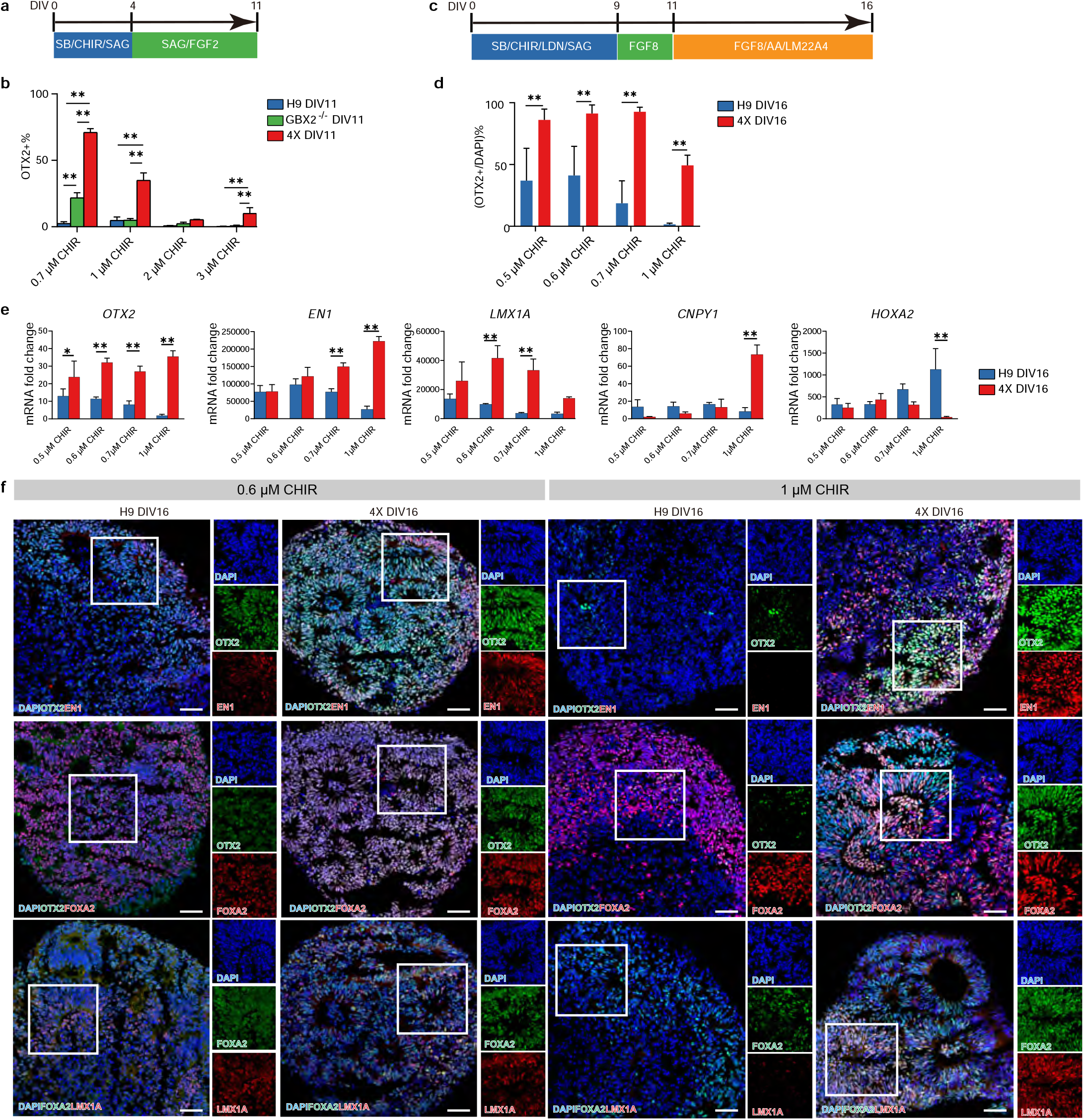
Differentiation into ventral midbrain progenitors. **a**, Schematic diagram of the 11 DIV CNP differentiation protocol using different concentrations of GSK3i ranging from 0.7 µM to 3 µM. **b**, Flow cytometry analysis of the percentage of OTX2-positive cells among H9, GBX2^-/-^ and 4X CNPs at 11 DIV. The data are presented as the mean ± SD; n= 3. Two-way ANOVA followed by Tukey’s multiple comparisons test comparing groups treated with the same concentration of GSK3i. **P< 0.01. **c**, Schematic diagram of the 16 DIV caudal midbrain differentiation protocol using different concentrations of GSK3i ranging from 0.5 µM to 1 µM. **d**, Quantification of OTX2-positive cells among H9 and 4X cells at 16 DIV after administration of GSK3i at concentrations ranging from 0.5 µM to 1 µM. The data are presented as the mean ± SD; n= 4-8. Two-way ANOVA followed by Sidak’s multiple comparisons test comparing groups treated with the same concentration of GSK3i. **P< 0.01. **e**, Expression of *OTX2, EN1, LMX1A, CNPY1*, and *HOXA2* in H9 and 4X cells treated with a range of GSK3i concentrations at 16 DIV. The data are presented as the mean ± SD., n= 3. Two-way ANOVA followed by Sidak’s multiple comparisons test comparing groups treated with the same concentration of GSK3i. *P< 0.05; **P< 0.01. **f**, Representative immunohistochemical analysis of OTX2, EN1, FOXA2, and LMX1A expression in H9 and 4X cells treated with GSK3i at concentrations of 0.6 µM and 1 µM on 16 DIV. DAPI was used as a nuclear stain. Scale bars, 50 µm. Caudal neural progenitor: CNP.

Our main objective was to generate LR-USCs that more efficiently generate mesDA neurons, even under suboptimal conditions. Thus, we compared H9 and 4X cells and differentiated them using a recently reported mesDA protocol (Fig. 2c) ^14^. This protocol is known to require adjustments to the concentration of GSK3i between cell lines; therefore, we started by titrating GSK3i from a concentration of 0.5 µM to 1 µM to identify the optimal concentration for generating posterior midbrain cells from H9 cells. First, we determined that the highest percentage of OTX2-positive H9 cells was obtained with a concentration of GSK3i between 0.5 and 0.7 µM and that there was a significant decrease in the number of these cells when the GSK3i concentration reached 1 µM (Fig. 2d). Second, we examined the expression levels of midbrain genes (Fig. 2e). The transcript level of *OTX2* in H9 cells at 16 DIV reached a maximum at a GSK3i concentration of between 0.5 and 0.6 µM, and *EN1* expression was the highest at a GSK3i concentration of 0.6 µM. Furthermore, the transcript level of the hindbrain gene *HOXA2* reached the lowest point at a GSK3i concentration between 0.5 and 0.6 µM. These results were consistent with previous reports, which indicated that a GSK3i concentrations below 1 µM is necessary for midbrain specification and that concentrations approaching 1 µM or higher result in a dramatic shift to hindbrain identity ^11^. Interestingly, *HOXA2* was expressed under optimal conditions, demonstrating that the protocol resulted in the production of a wide variety of cell types along the A-P axis, including hindbrain cell types, as reported by others ^11^. Overall, we determined that the optimal GSK3i concentration for H9 cells at 16 DIV was 0.6 µM (Fig. 2d, e and Supplementary Fig. 3).

We next compared H9 cells and 4X cells across multiple GSK3i concentrations and observed a significant improvement in the ability of 4X cells to produce midbrain cells. 4X cells produced a significantly higher percentage of OTX2-positive cells than H9 cells across all GSK3i concentrations (H9: 37.0%, 4X: 86.2%, P<0.0001 at 0.5 µM; H9: 41.1%, 4X: 91.3%, P<0.0001 at 0.6 µM; H9: 18.6%, 4X: 92.7%, P<0.0001 at 0.7 µM; H9: 1.5%, 4X: 49.3%, P<0.0001 at 1 µM; Fig. 2d). Furthermore, 4X cells exhibited significantly higher *OTX2* transcript levels than H9 cells across all GSK3i concentrations tested (P = 0.02 at 0.5 µM, P < 0.0001 at 0.6 µM, P = 0.0001 at 0.7 µM and P < 0.0001 at 1 µM; Fig. 2e) and *LMX1A* transcript levels are significantly higher than H9 cells at 0.6 and 0.7 µM GSK3i (P < 0.0001 at 0.6 µM and P = 0.0001 at 0.7 µM; Fig. 2e). In addition, we assessed the expression of caudal midbrain markers and found that *EN1* expression was significantly higher in 4X cells than H9 cells at 0.7 µM and 1 µM GSK3i (LogFC = H9: 77198.6, 4X: 149418, P = 0.0002 at 0.7 µM and H9: 27018.2, 4X: 222563, P<0.0001 at 1 µM) and that *CNPY1* expression was significantly higher in 4X cells than H9 cells at 1 µM GSK3i (LogFC = H9: 8.3, 4X: 73.4, P < 0.0001). Immunofluorescence staining confirmed the above results and showed that there was an abundance of EN1-positive 4X cells at GSK3i concentrations from 0.6 µM to 1 µM, whereas EN1-positive H9 cells were prominent only at a GSK3i concentration of 0.6 µM (Fig. 2f and Supplementary Fig. 3). These results demonstrated that 4X cells were more capable of generating caudal midbrain cells than H9 cells, even when exposed to concentrations of GSK3i that usually lead to the generation of hindbrain cell types.

### LR-USCs differentiated under conditions that favor a hindbrain identity generate midbrain progenitors

Using single-cell sequencing, we determined the cell types that were produced by H9 and 4X cells following the mesDA neuron differentiation protocol using 1 µM of GSK3i. Dimension reduction was performed by uniform manifold approximation and projection (UMAP), and there was a noticeable separation between 4X and H9 cells across clusters (Chi-square, P < 0.0001) and a difference in the cell types produced by the two cell lines at 16 DIV (Fig. 3a, d). This separation coincided with a marked shift in distribution along the A-P axis. Based on the expression of *OTX2* and *EN1*, we divided the A-P axis into four domains (rostral, *OTX2*-positive/*EN1*-negative; caudal midbrain, *OTX2*-positive/*EN1*-positive; rhombomere 1, *OTX2*-negative/*EN1*-positive; and posterior, *OTX2*-negative/*EN1*-negative) (Fig. 3c). 4X cells produced all four populations, with the smallest being the most rostral population (Fig. 3c). The caudal midbrain is the region where DA neurons of the substantia nigra develop, and 35.2% percent of 4X cells could be assigned to this region. Unlike for 4X cells, 0.6% of the cells produced by H9 expressing caudal midbrain markers and 97.5% of the cells were of the posterior population (*OTX2-*negative*/EN1-*negative; Fig. 3c, d). Overall, 4X cells preferentially generated *EN1*-positive cells (54.3%) spanning the midbrain and hindbrain, whereas the majority (97.5%) of H9 cells were classified as hindbrain cell types. Analysis of the ventral patterning gene *FOXA2* showed that it was highly expressed in both H9 and 4X cells, indicating that both cell lines were efficiently ventralized to a floor plate identity (Fig. 3d).

**Figure 3:**
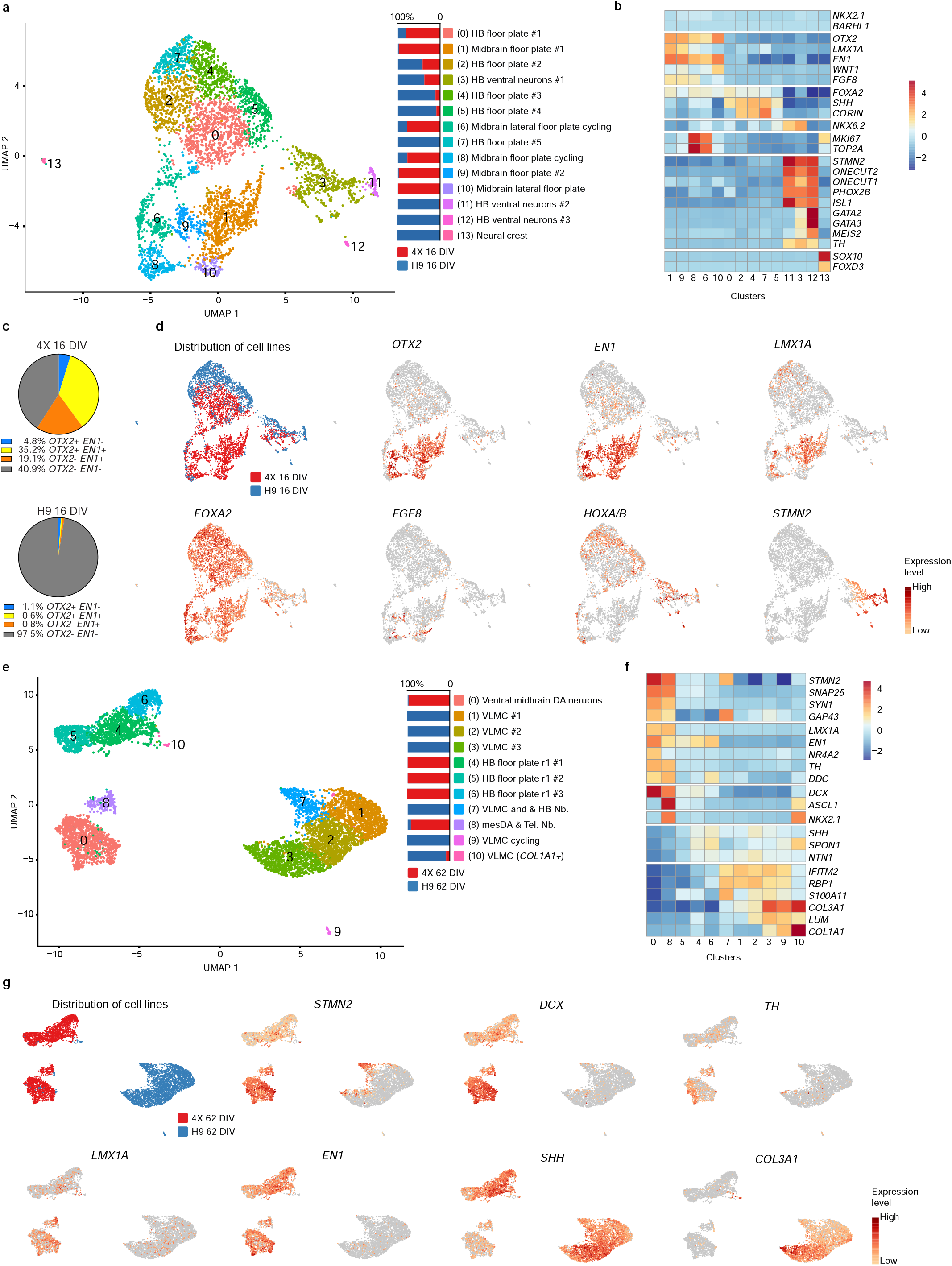
Single-cell sequencing of midbrain neurons differentiated using 1 µM GSK3i at 16 and 62 DIV. **a**, UMAP of cultured 4X and H9 cells at 16 DIV and a graph of the percentage of cells that each cell line contributed to each cluster (n = 10 4X cell spheres; n = 10 H9 cell spheres; total of 4,682 cells). **b**, Heatmap of the expression of selected genes illustrating the identity of the clusters. c, Percentage of 4X and H9 cells expressing *OTX2* and *EN1*. **d**, Feature plot of the contribution of each cell line to each cluster and feature plot of gene expression levels of *OTX2, EN1, LMX1A, FOXA2, FGF8, HOXA/B* family members and *STMN2*. **e**, UMAP of cultured 4X and H9 cells at 62 DIV and a graph of the percentage of cells that each cell line contributed to each cluster (n = 4 4X cell cultures; n = 4 H9 cell cultures; total of 6,804 cells). **f**, Heatmap of the expression of selected genes illustrating the identity of the clusters. **g**, Feature plot of the contribution of each cell line to each cluster and feature plot of the gene expression levels of *STMN2, DCX, TH, LMX1A, EN1, SHH* and *COL3A1*.

Upon further examination of caudal midbrain cells by graph-based clustering, we identified three clusters (clusters 1, 9, and 8) enriched in caudal midbrain floor plate progenitors expressing *FOXA2, OTX2, LMX1A* and *EN1* (Fig. 3a-d). Two additional midbrain floor plate clusters (clusters 6 and 10) were identified; these populations expressed *FOXA2, OTX2* and *EN1* but lacked *LMX1A*, indicating that they were a lateral floor plate population (Supplementary Fig. 4a). Clusters 8 and 6 were in a proliferative state, as revealed by the expression of *MKI67* and *TOP2A* (Fig. 3b). Cluster 1 was the largest midbrain population and exhibited the highest expression of midbrain markers, with 4X cells making up 98.6% of cells in this cluster.

By analyzing the hindbrain cells in more detail, we identified five clusters that we classified as hindbrain floor plate progenitors (clusters 0, 4, 2, 7, and 5), which expressed *FOXA2, SHH* and *CORIN* (Fig. 3b). The hindbrain floor plate clusters comprised both H9 and 4X cells (Fig. 3c). Further examination of *HOX* gene expression within these clusters showed that there was an abundance of cells expressing anterior *HOXA/B* genes (Fig. 3d). A total of 92.2% of the *HOXA/B* cells originated from H9 cells, confirming that H9 cells were of a more caudal identity than 4X cells (Fig. 3d and Supplementary Fig. 4b-c). We also identified a small population of early neural crest progenitors expressing *SO×10* and *FOXD3* (cluster 13), which were exclusively H9 cells (Fig. 3a, b).

By 16 DIV, three neuronal clusters (3, 11, and 12), which were predominately derived from H9 cells, were present (H9: 52%, 95%, 100%; 4X: 48%, 5%, 0%, respectively). Clusters 11 and 3 expressed high levels of *ONECUT2, PHOX2B* and *ISL1*, which are markers of early-born basal-plate hindbrain motor neurons ^35–37^. The remaining cluster, cluster 12, contained only 28 cells that expressed high levels of *GATA2, GATA3* and *MEIS2* but did not express *GAD1* or *GAD2*, indicating that they were immature V2b GABAergic neuroblasts ^36^. A population of cells within the neuronal clusters expressed tyrosine hydroxylase (*TH*); however, after subclustering, we found that these cells did not express *NR4A2* (also known as *NURR1*), *LMX1A* or *EN1*, indicating that they were not of a mesDA identity (Fig. 3b, d and Supplementary Fig. 4d, e).

### LR-USCs efficiently generate mesDA neurons under conditions that favor a hindbrain identity

To examine the extent to which midbrain floor plate progenitors can produce mesDA neurons, we extended the 1 µM GSK3i protocol to 62 DIV. At 62 DIV, the two cell lines occupied almost completely separate clusters (Chi-square, P < 0.0001; Fig. 3e, g). 4X cells were broadly be divided into two main cell types: hindbrain r1 floor plate clusters expressing *FOXA2, SHH, NETRIN1, SPON1 and EN1* (clusters 4, 5 and 6)^38^ and neuronal clusters (clusters 0 and 8; Fig. 3e, f). The neuronal clusters contained mesDA neurons identified by the expression of *TH, FOXA2, LMX1A* and *EN1* (Fig. 3f and Supplementary Fig. 4g). Clusters 0 and 8 was comprised almost entirely of 4X cells (4X: 82% and 77%, H9: 18% and 23%; Fig. 3e). Upon closer inspection of the difference between clusters 0 and 8, we identified a subset of cells within cluster 8 that expressed *NKX2*.*1*, a marker of hypothalamic neurons ^39^.

In contrast to 4X cells, H9 cells formed one main connected set of clusters (clusters 1, 2, 3, and 7) and two small isolated clusters (clusters 9 and 10; Fig 3e). All six clusters were dominated by cells expressing markers indicative of vascular leptomeningeal cells (VLMCs), i.e., *COL3A1, IFITM2* and *S100A11* (Fig. 3f). Interestingly, *EN1* was largely absent from the VLMC clusters (Fig. 3g). Cluster 7 also contained a population (41%) of cells expressing *STMN2, SEMA3C* and *PDLIM1* (Fig. 3f, Supplementary Fig. 4f), which, according to a single-cell brain atlas, corresponded to a subtype of peripheral sensory neurons ^40^.

We next wanted to examine the subtypes of mesDA neurons by first subclustering of *TH*-positive neurons (Supplementary Fig. 4h-j). To distinguish between substantia nigra and ventral tegmental area (VTA) DA neurons, we assessed the expression of *GIRK2* (also known as *KCNJ6*) and CalbindinD (*CALB1*). *GIRK2* was highly expressed in the subclusters 1 and 3, and accounted for 38% of the TH population, and only a small proportion of *TH* neurons expressed *CALB1* (13.6%; Supplementary Fig. 4k).

To further support our single-cell sequencing results, histological analysis was performed. We used the same growth factor paradigm but adapted a differentiation protocol to generate organoids to provide an optimal environment for the survival of neurons (Fig. 4a). At 83 DIV, we observed a large population of FOXA2/TH double-positive DA neurons within organoids produced from 4X cells (Fig. 4b). In contrast, TH-positive cells were occasionally scattered throughout organoids produced by H9 cells; however, we rarely detected TH-positive cells coexpressing FOXA2 (Fig. 4b). These results were consistent with the histological analysis of our 2D cultures at 62 DIV (Supplementary Fig. 5). Further examination of TH-positive neurons showed that, in accordance with our single-cell data, the most abundant population of TH neurons derived from 4X cells coexpressed GIRK2 (Fig. 4b) and that there was a small population of CALB1/TH double-positive neurons (Fig. 4b).

**Figure 4:**
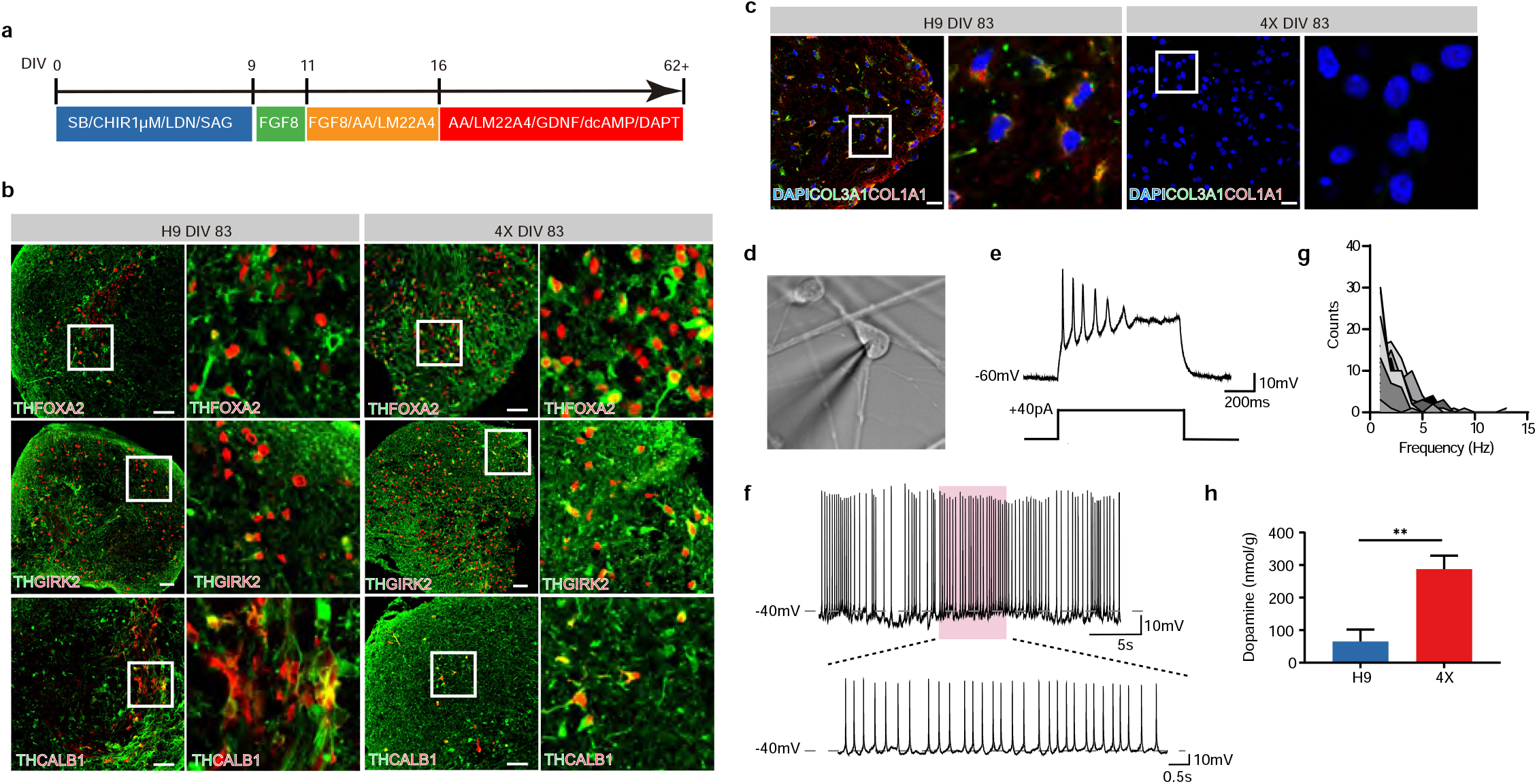
Generation of functional ventral midbrain DA neurons in vitro. **a**, Schematic diagram of the long-term (62 DIV) neuronal differentiation protocol. **b**, Representative immunohistochemical images of TH/FOXA2, TH/GIRK2, and TH/CALB1 costaining in H9 and 4X cells treated with 1 µM GSK3i on 83 DIV. Scale bars, 50 µm. **c**, Representative immunohistochemical analysis of COL3A1 and COL1A1 expression. Many H9 cells were double positive for COL3A1 and COL1A1, but no 4X cells were positive for COL3A1 or COL1A1. DAPI was used as a nuclear stain. Scale bars, 20 µm. **d**, Phase contrast image of a patched 4X neuron during whole-cell recording. Scale bar, 10 µm. **e**, Representative response (top trace) to a depolarizing current injection (bottom trace) showing firing of repetitive action potentials. **f**, Example of spontaneous firing at a resting membrane potential of -45 mV showing burst-like events. Overshooting spikes occurred in groups interspersed by periods of subthreshold membrane oscillation. **g**, Frequency distribution of spontaneous cell firing showing firing frequencies ranging between 1 and 5 Hz (n = 16 cells). **h**, Dopamine content (normalized to the protein concentration) in 4X and H9 cells at 79 DIV, as measured by HPLC. The data are presented as the mean ± SD; n= 3. An unpaired t-test was used to compare groups. **P< 0.01.

According to the single-cell sequencing data, the majority of H9 cells were VLMCs (Clusters 1, 2, 3, 7, 9, 10; 93% of H9 cell). To confirm this finding, we examined the expression of VLMC markers in organoids at 83 DIV, and we identified a large population of COL3A1/COL1A1 double-positive cells with a nonneuronal morphology among H9 cells (Fig. 4c). No cells positive for COL3A1 or COL1A1 were identified among 4X cells (Fig. 4c).

### DA neurons derived from LR-USCs exhibit pacemaker activity

Having shown that we can generate mesDA neurons from 4X cells under caudalizing conditions, we next wanted to examine the electrophysiological properties of the DA neurons. We performed *in vitro* electrophysiological recordings in whole-cell patch-clamp configuration between DIV 80 and DIV 84 (Fig. 4d). We observed that the cells developed into electrophysiologically mature neurons, as measured by their ability to generate repetitive action potentials upon somatic current injection (Fig. 4e). Recordings in current-clamp mode revealed spontaneous pacemaker activity characteristic of a DA neuron identity, with a mix of single spikes and phasic bursts (Fig. 4f). Membrane oscillations collapsed at potentials below -50 mV (data not shown). The firing frequency in our sample ranged from 1 to 5 Hz (Fig. 4g). Furthermore, HPLC analysis of cell extracts showed that the DA content in the 4X cells was significantly higher than that in the H9 cells (287.4 nmol/g in 4X cells vs. 65.1 nmol/g in H9 cells, P = 0.002; Fig. 4h).

### Analysis of 4X cells *in vivo* in a Parkinson’s disease rat model

When using current DA neuron differentiation protocols, DA neurons account for only a small percentage of cells of the entire graft when mesDA progenitors are grafted *in vivo*. A protocol using FGF8 to induce midbrain caudalization was shown to result in the production of approximately 3,700 TH-positive DA cells per 100,000 grafted cells ^14^, and a more recently reported protocol using a delayed boost of WNT was found to induce the generation of 9,173 TH-positive cells per 6.22 mm^3^ graft following transplantation of 450,000 cells ^13^. We investigated how our 4X LR-USCs behave *in vivo* when transplanted into a rodent model of Parkinson’s disease. To confirm the results of our single-cell analysis, we used the same midbrain differentiation protocol with an unfavorable caudalizing concentration of GSK3i (1 µM).

A total of 250,000 4X cells or H9 cells were transplanted into the striata of nude rats with 6-OHDA-induced medial forebrain bundle (MFB) lesions 4 weeks after lesioning. A third group of lesioned rats that did not undergo transplantation was used as a lesion control (6-OHDA, see Fig. 5a for study design). At the time of transplantation, all three groups of rats exhibited a similar number of amphetamine-induced ipsilateral rotations/min (limit for inclusion: 5 rotations/min, Supplementary Fig. 6a), confirming significant loss of DA striatal innervation. All three groups showed forelimb asymmetry in the cylinder test, with the rats using mostly the ipsilateral forepaw (6-OHDA 52.4%; H9 70.4% and 4X 67.4% of total) and almost never the contralateral forepaw (6-OHDA 1.3 %; H9 1.3% and 4X 0% of total) to touch the walls or land on the floor after rearing, further supporting the induction of a DA deficit by 6-OHDA (Supplementary Fig. 6b).

**Figure 5:**
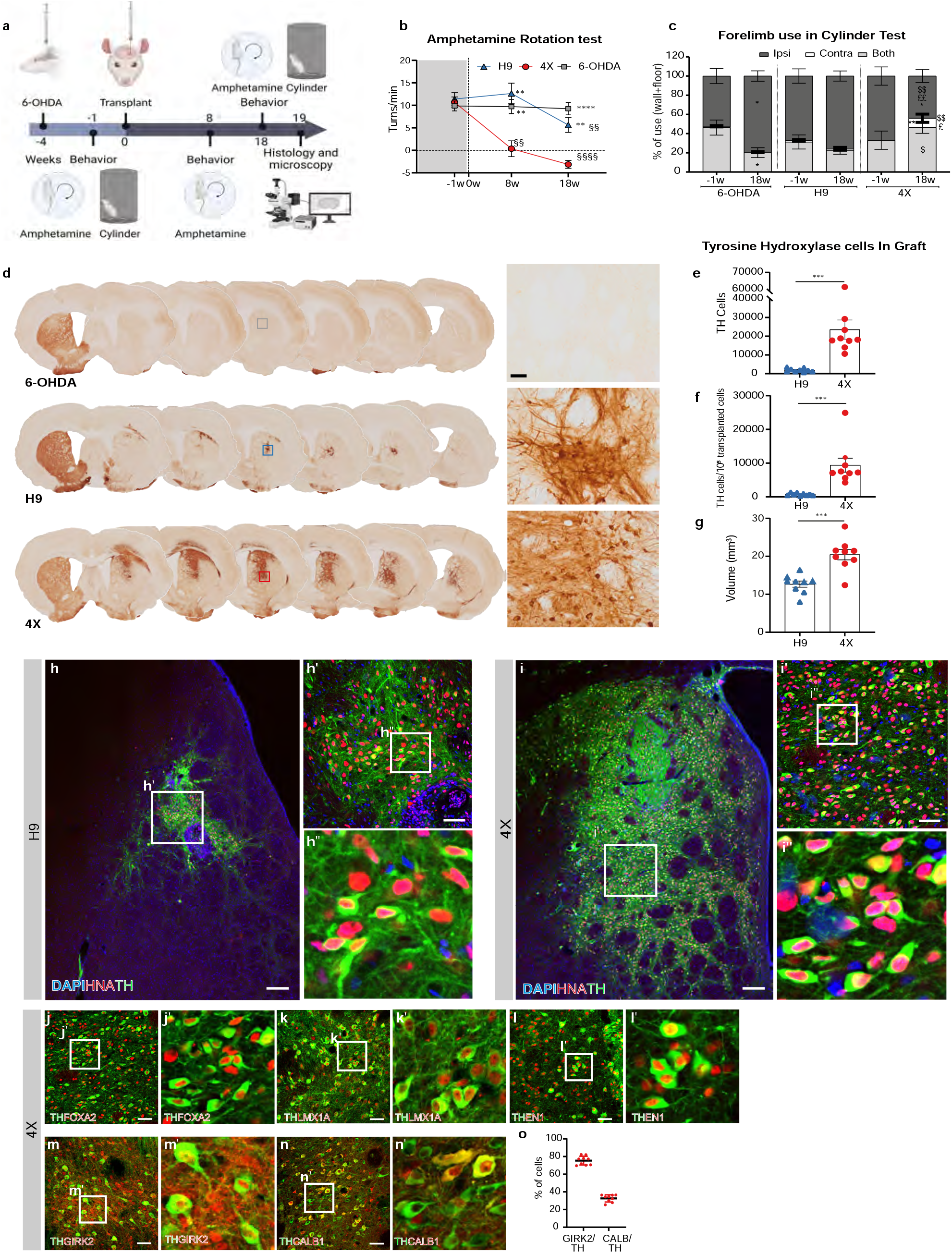
In vivo analysis of cells transplanted into a Parkinson’s disease rat model. **a**, Overview of the in vivo study. Unilateral 6-OHDA-induced MFB lesions were generated (week -4) and confirmed 3 weeks later by the cylinder and amphetamine-induced rotation tests. The animals were subdivided into 3 groups with similar average scores on the rotation test. Four weeks after lesioning (week 0), two of these subgroups were transplanted with 250,000 cells (H9 or 4X cells), and the third group did not undergo transplantation (6-OHDA lesion group). The rotation test was repeated at weeks 8 and 18 posttransplant, and the cylinder test was repeated at week 18. The animals were killed at week 19 posttransplantation (23 week after lesioning) for histological analysis. **b**, Amphetamine-induced rotational asymmetry. Two-way repeated measures ANOVA followed by Sidak’s multiple comparison test; time: F(1.689, 35.46) =19.50, P<0.0001; treatment: F (2, 21) = 15.23 P< 0.0001. ** P< 0.01 and ****P< 0.0001 vs. the 4X cell-transplanted group at the same time point. §§ P< 0.01 and §§§§P<0.0001 vs. the same group at week -1. **c**, The use of each forelimb (contra or ipsi) and both forelimbs in the cylinder test was analyzed by two-way repeated measures ANOVA followed by Sidak’s multiple comparison test with time and group as variables. Time x group: both: F (2, 22) = 5.785, P=0.009; ipsi: F (2, 22) = 8.800, P=0.001; contra: F (2, 22) = 4.642, P=0.021. *P<0.05 and **P<0.01 vs. the same group at -1 week. $P<0.05 and $$P<0.01 vs. the 6-OHDA lesion group at the same time point. £P<0.05 and ££P<0.01 vs. the H9 cell-transplanted group at the same time point. The data in **(b)** and **(c)** are presented as the mean ± SEM. n= 7 rats in the 6-OHDA lesion group, n = 9 rats in the 4X cell-transplanted group, and n=8 rats in the H9 cell-transplanted group). **d**, Representative photos of coronal sections from all three groups immunostained for TH. Higher magnification images of the areas in the frame are shown on the right. Scale bars, 50 µm for all three photos in the column. The graphs on the right show **(e)** the estimated numbers of TH-positive cells in the grafts, **(f)** the yield of TH-positive neurons per 100,000 grafted cells and **(g)** the volume of the TH-positive graft (see Methods for details). **h-i**, Representative photomicrographs showing HNA-positive and TH-positive cells within H9 cell **(h)** and 4X cell **(i)** grafts. The squares in **h-i** and **h’-i’** indicate the magnified areas shown in **h’-i’** and **h’’-i’’**, respectively. DAPI was used as a nuclear stain. Scale bars, 200 μm **(h-i)** and 50 μm (**h’-i’)**. **j-n**, Representative immunofluorescence images of cells double-positive for TH/FOXA2 **(j)**, TH/LMX1A **(k)**, TH/EN1 **(l)**, TH/GIRK2 **(m)** and TH/CALB1 **(n)** within 4X cell grafts. **(j’-n’)** High-power images of j-n highlighting the graft composition. Scale bar, 50 μm. **o**, Quantitative analysis of the immunofluorescence data in m and n, showing the percentages of GIRK2/TH and CALB1/TH double-positive cells within 4X cell grafts. The data are presented as the mean percentage ± SD (n=9 rats).

Eight weeks posttransplantation, rats that received 4X cells showed complete correction of amphetamine-induced ipsilateral rotation (pretransplant: 10.6 vs 8w: 0.35 rotations/min), suggesting that a sufficient amount of dopamine was released in the striatum to normalize (Fig. 5b) or even overcompensate for this behavior, as suggested by the number of contralateral rotations (−3.12 rotations/min) observed 18 weeks posttransplantation. However, H9 cells-transplanted rats presented a statistically similar number of ipsilateral rotations as the control 6-OHDA lesion group (pretransplant: 9.8; 8w: 9.7 and 18w: 9. 2 rotations/min) throughout the entire experiment, showing only a significant reduction in the number of rotations compared to pretransplantation values at 18 weeks (H9 pretransplant: 11.5; 8w: 12.6 and 18w: 5.6 rotations/min, Fig. 5b). Analysis of spontaneous motor behavior in the cylinder test confirmed the significant improvement in 4X cell-transplanted rats, as these rats used the contralateral forelimb alone (9.7% of total) or together with the ipsilateral forelimb (both 46.3%) during the test at week 18 (Fig. 5c). However, both H9 cell-transplanted and 6-OHDA lesion rats used mostly the ipsilateral forelimb (76.1% and 79.3% of total respectively), using both forelimbs less than 30% of the time but almost never using the contralateral impaired forelimb when rearing in the cylinder, as observed before transplantation (Fig. 5c). Therefore, 4X cell transplantation significantly improved both drug-induced and spontaneous motor behavior after 6-OHDA-induced lesioning of the MFB.

Postmortem histological analysis of the brains showed that rats transplanted with 4X cells had graft-derived TH-positive cells in the area of injection, i.e., the striatum, as well as in globus pallidus, the corpus callosum and the area of the cortex above the striatum (Fig. 5d). However, H9-derived TH-positive cells remained mostly in the striatum and were also found in the globus pallidus in few animals. Quantification of graft-derived TH-positive cells (in the striatum and globus pallidus) showed that there were significantly more TH-positive cells per graft in 4X cell-transplanted rats (an average of 23,520 TH-positive cells per graft) than H9 cell-transplanted rats (1,898 TH-positive cells per graft), resulting in a larger yield (9,408 TH-positive cells per 100,000 transplanted 4X cells vs. 759 TH-positive cells per 100,000 transplanted H9 cells) (Fig. 5e, f). The TH-positive 4X cell graft extended across 6-8 coronal A-P striatal sections (in a series of 8), while the H9 cell graft occupied 4-5 sections. Thus, the estimated graft volume was 61% larger in the 4X cell-transplanted rats (20.46 mm^3^) than in the H9 cell-transplanted rats (12.69 mm^3^) (Fig. 5g). The increase in TH-positive cell number resulted in a significantly higher density of TH cells in the graft in the 4X cell-transplanted group (1,090 ± 464 cells/mm^3^ vs. 143 ± 49 cells/mm^3^ in the H9 cell-transplanted group; P<0.0001), which is in agreement with the rapid and robust behavioral recovery observed in the 4X cell-transplanted group.

Further examination of the graft showed that all TH-positive neurons identified within the 4X and H9 cell grafts coexpressed the human nuclear marker human nuclear antigen (HNA) (Fig. 5h, i). In the grafts of 4X cell-transplanted rats, TH-positive neurons coexpressed FOXA2, LMX1A and EN1, indicating that they were mesDA neurons (Fig. 5j-l). To distinguish between A9 and A10 neurons, we calculated the proportion of TH-positive neurons expressing GIRK2 and CALB1 and found that 75.4% ± 4.99 of TH-positive neurons were GIRK2-positive (Fig. 5m-o).

Interestingly, we also found that TH-positive neurons derived from H9 cells were positive for FOXA2, LMX1A and EN1 (Supplementary Fig. 6c-e). This was in contrast to our *in vitro* experiments, in which TH-positive neurons derived from H9 cells rarely expressed FOXA2 (Fig. 4b, Supplementary Fig. 5a), suggesting that the *in vivo* environment is more permissive for the development and survival of TH-positive neurons than an *in vitro* environment. Since the *in vitro* data showed that H9 cells produced a large number of VLMCs, we examined the expression of vascular markers in the grafts of both 4X cell-transplanted rats and H9 cell-transplanted rats. Within the H9 cell grafts, we identified a large population of COL3A1/COL1A1/HNA triple-positive cells, whereas in 4X cell grafts, we rarely detected COL1A1-positive cells coexpressing the marker HNA (Supplementary Fig. 6f-g). Overall, the *in vivo* histological data showed that 4X cells were capable of producing a robust population of mesDA neurons, consistent with the rapid motor recovery seen at 8 weeks.

## Discussion

In this study, we engineered a novel type of stem cells, LR-USCs, with restricted differentiation potential. By knocking out genes involved in early lineage specification, we aimed to prevent the cells from differentiating into unwanted lineages to guide their differentiation toward the cell type of interest: mesDA neurons. Specifically, we examined the genes involved in patterning of the A-P axis because of the difficulties in fine-tuning differentiation to reproducibly generate pure caudal midbrain progenitors. We targeted transcription factors that are involved in the early specification of the hindbrain (GBX2) and spinal cord (CDX1/2/4). Importantly, the genes we targeted are not involved in the development of mesDA neurons. *CDX1/2/4* are expressed in the posterior region of the developing neural tube, which develops distinctly from the anterior neuroectoderm, and *GBX2* is initially expressed throughout the length of the hindbrain and spinal cord and is then restricted to the r1-3 regions. By knocking out these genes, we generated 4X LR-USCs, which efficiently generated midbrain cell types when differentiated under optimal or unfavorable (highly caudalizing) conditions.

Under caudalizing conditions, the majority of H9 cells adopted a floor plate hindbrain identity and produced a large population of nonneuronal cells expressing VLMC markers and only a small population of DA neurons. In contrast, when 4X cells were differentiated under the same conditions, a large population of ventral midbrain progenitors, which developed into DA neurons, was produced. Electrophysiological analysis of DA neurons showed that they were functional and displayed characteristic pacemaker activity. When 4X cells were transplanted *in vivo* into rats with 6-OHDA-induced MFB lesions, motor behavior improved, with amphetamine-induced rotation being fully corrected after only 8 weeks posttransplantation and the spontaneous use of the affected forelimb showing recovery after 18 weeks. In comparison, transplantation of cells subjected to differentiation protocols has been reported to achieve similar normalization of amphetamine-induced rotation after five months ^13,14^.

Histological examination of 4X cell grafts showed an estimated number of 23,520 TH-positive cells after 250,000 cells transplanted, which is substantially greater than that reported for other methods. This large number of TH-positive cells, along with the fact that the cells expressed FOXA2, EN1 and LMX1A, supports the observed rapid behavioral recovery. Furthermore, the majority of cells were GIRK2-positive, demonstrating that there was an abundance of DA substantia nigra neurons.

Thus far, we have described how knockout of four selected genes can dramatically increase the specification of PSCs to mesDA neurons by restricting the cell types along the A-P axis that they can differentiate into. It is possible to further restrict cell fate by knocking out additional genes to prevent differentiation into remaining populations of unwanted cells, which would further enhance the ability to generate mesDA neurons. Specifically, our single-cell sequencing data showed that 4X cells are capable of producing telencephalic and anterior hindbrain cell types. By targeting transcriptional determinates of these populations, we speculate that we could eliminate these populations. Furthermore, we can also target dorsal populations in addition to populations along the A-P axis. It is conceivable that by restricting the genome even further, we can produce a cell line that is capable of producing a highly pure population of mesDA neurons.

One of the important characteristics of the 4X cells is their ability to generate DA neurons under a broader range of growth factor conditions than other cells. This has significant advantages for clinical applications, allows for easier upscaling and reproducibility and reduces cell line variability. LR-USCs are not restricted to producing mesDA neurons; through deletion of other sets of genes, LR-USCs can be designed to generate other neural populations or cell types from other germ layers, which can be used for cell transplantation therapy or drug discovery for the treatment of a range of disorders.

## Methods

### hPSC culture

hESCs (H9 cell line, WiCell) were maintained on irradiated human fibroblasts in KSR medium consisting of DMEM/nutrient mixture F-12 supplemented with nonessential amino acids (NEAAs) 1%, glutamine 2 mM, 0.1 mM β-mercaptoethanol, 0.5% pen/strep and 20% knockout serum replacement. The KSR medium was supplemented with FGF2 (15 ng/ml; Peprotech) and Activin A (15 ng/ml; R&D Systems). Every seven days, the cells were manually passaged, and fragments were transferred to a freshly prepared gelatin-coated dish containing irradiated fibroblasts ^41^.

### Differentiation into CNPs

hESCs were differentiated into CNPs as described previously ^29^. Briefly, hESC fragments were cut from colonies growing on irradiated feeders (CCD-1079Sk, ATCC) and plated in vitronectin-coated plates in N2B27 medium containing neurobasal medium (NBM) and DMEM/F-12 supplemented with 1% N2 supplement at a 1:1 ratio, 1% B27 supplement minus vitamin A, 1% insulin/transferrin/selenium-A (ITS-A), 0.3% glucose, 1% Glutamax supplement, and 0.5% penicillin/streptomycin (all from Life Technologies). The medium was supplemented with SB431542 (SB; 10 μM, Tocris Bioscience) and CHIR99021 (CHIR; 3 μM, Stemgent) for 4 days. For the 11 DIV CNP differentiation protocol, cells were cultured as described above and supplemented with 400 nM SAG (Millipore). After day 4 the colonies were dissected into 0.5 mm pieces and cultured in suspension in low-attachment 96-well plates (Corning) in N2B27 medium supplemented with FGF2 (20 ng/ml; PeproTech) and 400 nM SAG.

### Differentiation into mesDA neurons

mesDA neurons were generated by implementing previously described protocols with minor modifications ^14^. Briefly, from day 0 to day 9, cells were grown in N2B27 medium supplemented with 10 µM SB, CHIR (0.5 to 1 µM), 0.1 µM LDN193189 (LDN; Stemgent), and 400 nM SAG (Millipore). On day 4, the colonies were cut into fragments and cultured in suspension. From day 9 to day 11, the supplements in the medium were replaced with FGF8 (100 ng/ml; R&D Systems). From day 11, the medium was supplemented with FGF8 (100 ng/ml), LM22A4 (2 µM), and ascorbic acid (200 µM; Sigma). On day 16, the cells were dissociated with Accutase and subsequently grown on culture plates coated with polyornithine, fibronectin, and laminin (all from Sigma). Neural differentiation medium consisting of 1% B27 supplement, 25 U/mL pen/strep, 0.5% Glutamax was supplemented with 200 µM ascorbic acid, LM22A4 (2 µM), 1 µM DAPT (Tocris Bioscience), GDNF (10 ng/ml), and dcAMP (500 µM). The medium was changed every second day until the end of the experiment. Alternatively, on day 16, the cultured cells were maintained in suspension to generate organoids.

### Generation of CRISPR lentiviral vectors

A pLV-4gRNA-GBX2-RFP lentiviral plasmid containing four CRISPR target sites in GBX2 was generated using the multiplex CRISPR lentiviral vector system ^42^. First, oligos containing the 20 bp protospacer sequence against the four CRISPR target regions in *GBX2* (Supplementary Table 1) were cloned by BbsI digestion and ligated into the following entry plasmids, i.e., ph7SK-gRNA, phU6-gRNA, pmU6-gRNA and phH1-gRNA (Addgene #53189, 53188, 53187 and 53186), to generate four gRNA GBX2 entry plasmids. Second, using the golden recombination method, the pLV-GG-hUbC-dsRED plasmid (Addgene # 84034) and the gRNA GBX2 entry plasmids were recombined by BsmBI digestion and ligation to form the final pLV-4gRNA-GBX2-RFP plasmid. A multiplex CRISPR plasmid containing CRISPR targets in the *GBX2* and *CDX1/2/4* genes was generated in a similar manner as above. Four entry plasmids, i.e., ph7SK-GBX2-gRNA, phU6-CDX4-gRNA, pmU6-CDX1-gRNA and phH1-CDX2-gRNA, were generated and recombined with pLV-hUbC-Cas9-T2A-GFP (Addgene #53190), resulting in the generation of the pLV-hUbC-GBX2-CDX124-Cas9-T2A-GFP plasmid (see Supplementary Table 1 for gRNA sequences).

### Generation of knockout cell lines

Three lentiviral plasmids, pLV-4gRNA-GBX2-RFP, pLV-hUbC-GBX2-CDX124-Cas9-T2A-GFP and lentiCas9-Blast (Addgene # 52962), were used to produce lentiviruses. Lentiviral production was performed as described previously ^43^. To generate the GBX2 knockout cell line, H9 cells were transduced with LV-4gRNA-GBX2-RFP and lentiCas9-Blast, and after three days, transduced cells were selected using 10 µg/ml blasticidin for 6 days (Supplementary Fig. 1). FACS was then used to separate single RFP-positive cells in a 96-well plate using the 561 nm laser on a FACSAriaIII (BD Biosciences, San Jose, CA). Indels at the corresponding target sites in the clones were analyzed by genomic PCR. To generate the 4X knockout cell line, H9 cells were infected with LV-hUbC-GBX2-CDX124-Cas9-T2A-GFP, and after 7 days, single GFP-positive cells were sorted by FACS (Supplementary Fig. 2). Allele-specific mutations in both the *GBX2*^-/-^ and 4X cell lines were confirmed using whole-exome sequencing. Whole-exome sequencing and mapping were performed by BGI (BGI, Copenhagen). Integrated Genome Browser V 2.10.0 was used to identify allele-specific mutations. To identify large deletions that could not be mapped by the alignment tools, individual sequencing reads were extracted from the FastQ files using Grep and manually analyzed.

### QPCR and NanoString

For QPCR and NanoString experiments, RNA was extracted using the Qiagen RNeasy mini kit and treated with DNase I according to a standard protocol. cDNA was generated from 500 ng of total RNA using Superscript III and random primers following the manufacturer’s instructions. For QPCR, TaqMan Universal Master mix II without UNG and TaqMan probes were used (Supplementary Table 2). NanoString experiments were performed using the NanoString nCounter SPRINT (NanoString Technologies) according to the manufacturer’s instructions. Briefly, 200 ng of total RNA was used. Reporter probes were hybridized for 20 hours at 65 °C. A custom designed NanoString CodeSet consisting of a panel of capture and reporter probes designed to target 100 nucleotides of the gene of interest and a panel of housekeeping genes was used (Supplementary Table 3). RNA expression data were normalized to the expression of housekeeping genes.

### Immunofluorescence

Cells cultured on glass coverslips or suspended in culture plates as spheroids were collected. The samples were washed with PBS two times, fixed in 4% paraformaldehyde (PFA) in PBS at 4 °C for 15 min (glass coverslips) or 2 hours (spheroids), and washed 3 times with PBS for 10 min each. The spheroids were transferred to 20% sucrose in PBS, incubated at 4 °C overnight and embedded in OCT (Tissue-Tek). Sections were cut at a thickness of 10 µm using a cryostat (Crostar NX70) at -20 °C. The coverslips and sections were incubated in 0.25% Triton X in PBS (PBST) for 10 min and blocked in 5% donkey serum (Almeco) in PBST for 1 hour at room temperature. The following primary antibodies were applied overnight at 4 °C: goat anti-OTX2 (1:500, R&D Systems, cat# AF1979), mouse anti-CDX2 (1:200, BioGenex, cat# MU392-UC), mouse anti-Engrailed1 (EN1, 1:40, DSHB, cat# 4G11-s), rabbit anti-EN1 (1:50, Merck, cat# HPA073141), rabbit anti-FOXA2 (1:500, Cell Signaling, cat# 8186), goat anti-FOXA2 (1:200, R&D Systems, cat# AF2400), rabbit anti-LMX1A (1:5000, Millipore, cat# AB10533), mouse anti-TH (1:2000, Millipore, cat# MAB318), rabbit anti-TH (1:1000, Pel Freez, cat# P40101-150), rabbit anti-GIRK2 (1:500, Alomone, cat# APC-006), mouse anti-CALB1 (1:5000, SWANT, cat #300), rabbit anti-Collagen3A1 (1:1000, NovusBio, cat# NB120-6580), sheep anti-hCOL1A1 (1:200, R&D Systems, cat# AF6220), and mouse anti-HNA (1:200, Abcam, cat# ab191181). After the cells were washed with PBST three times for 10 min each, corresponding secondary antibodies (1:200, Jackson ImmunoResearch Laboratories or 1:1000, Invitrogen) were applied for one hour at room temperature. After the secondary antibodies were removed, the cells were washed three times with PBST for 10 min each in the dark. The nuclei were counterstained with DAPI (1 μg/ml, Sigma) and rinsed with PBS three times for 5 min each. The slides or coverslips were mounted with PVA-DABCO. Images were captured with a confocal microscope (Zeiss LSM 780) and Zen software.

### Flow cytometry analysis

The cells were washed two times with PBS- and dissociated with Accutase to obtain single cells. The cells were centrifuged at 300×g for 4 min and resuspended in 4% PFA for 10 min at room temperature. Then, the cells were washed with PBS-, centrifuged, resuspended in PBST, centrifuged again, and blocked in 5% donkey serum for 30 min at room temperature. Primary antibodies in blocking solution were added to the cells, and the cells were incubated for 2 hours at room temperature. The cells were washed once with PBST, resuspended in secondary antibodies in blocking solution and incubated for 30 min at room temperature in the dark. The cells were washed with PBST overnight at 4 °C and resuspended in PBS for flow cytometry using a NovoCyte Quanteon analyzer (Acea Biosciences Inc., Santa Clara, CA). The data were analyzed with FlowJo software (v. 10, Ashland, OR).

### Quantification of immunofluorescence images

The percentages of OTX2/DAPI double-positive, GIRK2/TH double-positive and CALB1/TH double-positive cells, either in culture or within a graft, were quantified with ImageJ software (1.53) by semiautomatic object-based colocalization analysis ^44^. The Colocalization Image Creator Plugin was used to process the multichannel immunofluorescence images into multichannel binary and grayscale output images. Binary output images were generated by processing input channels for ImageJ filters that applied an automatic local intensity threshold, radius outlier removal, watershed segmentation, eroding, hole filling, Gaussian blurring and maximum algorithms. Binary objects of an inappropriately small size were further removed from the output images via a defined minimum area size. To improve the visualization of the colocalization signals, the object overlap was restricted to the nuclei of the cells. The accuracy of the binary object segmentation was visually verified via grayscale output images. Once verified, the binary objects, representing either individually labeled or colabeled cells, were quantified automatically using the Colocalization Object Counter plugin. All immunofluorescence images were analyzed with conserved binary object segmentation settings. A minimum of 4 random fields captured at 20x and 63x were used for the quantification of OTX2 positive cells in culture. Quantification of GIRK2/TH double-positive and CALB1/TH double-positive cells within the graft was performed blindly by analyzing 4 nonoverlapping images taken at 20x from 2 sections per graft per animal.

### RNA sequencing and data analysis

Library construction, sequencing and initial data filtering, including adaptor removal, were performed by the BGI Europe Genome Center. Total RNA was subjected to oligo dT-based mRNA enrichment. Sequencing of 100 bp paired-end reads was performed on the DNBseq platform. More than 20 million clean reads were obtained per sample. The reads were aligned to the Human genome build hg38 (Ensemble release 92) using HISAT2 aligner (v2.1.0) ^45^. Transcript quantification was performed using FeatureCount (v1.6.4), and the read counts were normalized for effective gene length and sequencing depth to yield transcripts per kilobase million (TPM) ^46^. Differentially expressed genes were identified from count tables using edgeR (v3.32) ^47^. Centering and univ variance scaling were applied to TPM values to construct heatmaps and perform principal component analysis (PCA) by Clustvis using SVD with imputation ^48^.

### Single-cell RNA-seq and data analysis

On days 16 and 62, cultured cells were dissociated into single cells using Accutase. On day 16, neurospheres (n = 10 biological replicates per cell line) were pooled together, and on day 62, four biological replicates per cell line were pooled together. To construct the library, the 10X Genomics Chromium Next GEM Single Cell 3’ kit v 3.1 was used according to a standard protocol. Each of the four groups (day 16 H9 cells, day 16 4X cells, day 62 H9 cells and day 62 4X cells) was run in separate lanes of the Chromium controller, and a total of 8,000 cells were loaded per lane. Next-generation sequencing was performed by the NGS Core Center, Department of Molecular Medicine, Aarhus University Hospital, Denmark. Sequencing was performed on an Illumina NovaSeq instrument. The Cell Ranger Single-Cell Software Suite (v 3.1.0) was used for sample demultiplexing, barcode processing and single-cell 3′ gene counting. The reads were aligned to the human GRCh38 reference genome. Further analysis, including quality filtering, dimensionality reduction, and application of standard unsupervised clustering algorithms, was performed using the Seurat R package (v 3.2.1). To exclude outlier cells, the number of genes expressed in each cell was plotted for each sample to select the optimal allowed minimum number of genes per cell. The minimum numbers of genes per cell were set to 3000 for day 16_H9 cells, 2000 for day 16_4X cells, 3000 for day 62_H9 cells, and 3000 for day 62_4X cells. Cells with a high percentage of reads mapped to mitochondrial genes were also removed. For day 16 samples, all cells with more than 10% mitochondrial RNA were removed; for day 62, the limit was 15%. The R package DoubletFinder (v.2.0.3) was used to remove cell doublets from the single-cell transcriptome data, with the expected percentage of doublet cells being set at 7.5%. The single-cell data were normalized by dividing the gene counts of each cell by the total counts for that cell, multiplying by a scaling factor of 10,000, and natural-log transforming the result. Dimensionality reduction was performed using the UMAP technique. Clustering was performed by Seurat’s graph-based clustering approach using the FindClusters function, with the resolution set to 0.6. Various single-cell plots were generated using Seurat in R.

### Electrophysiology

Electrophysiological recordings of 4X cells were performed at 80-84 DIV. 4X cells cocultured with astrocytes on 13 mm Ø coverslips were transferred to the recording chamber following progressive transition from culture medium to artificial cerebrospinal fluid (aCSF) by adding five drops (200 µL each) of aCSF to the cultured medium over 20 s. After being transferred into the recording chamber, the coverslips were continuously perfused at room temperature with aCSF containing (in mM) 119 NaCl, 2.5 KCl, 26 NaHCO_3_, 1 NaH2PO_4_, 11 D-glucose, 2 CaCl_2_, and 2 MgCl_2_ (adjusted to pH 7.4).

The recording chamber was mounted on an upright microscope (Scientifica) linked to a digital camera (QImaging Exi Aqua). The 4X cells were visualized using a 63X water-immersion objective (Olympus, LumiPlan). The cells selected for electrophysiological recordings exhibited a neuron-like morphology with fine branching neurites. Clusters of amassed cells were avoided. Acquisitions were performed in whole-cell configuration in current-clamp mode using Clampex 10.6 software connected to a Multiclamp 700B amplifier via a Digidata 1550A digitizer (Molecular Devices). The data were low-pass filtered at 200 Hz and digitized at 10 kHz, and the whole-cell capacitance was compensated. Patch pipettes (resistance of 5-10 MOhm) were filled with an internal solution containing (in mM) 153 K-gluconate, 10 HEPES, 4.5 NaCl, 9 KCl, 0.6 EGTA, 2 MgATP, and 0.3 NaGTP. The pH and osmolarity of the internal solution were close to physiological conditions (pH 7.4, osmolarity of 297 mOsm). The access resistance of the cells in our sample was ∼ 30 MOhm. Among the recordings of 30 neurons that were obtained, 16 were kept for analysis. The rest of the recordings were from neurons that either were nonrespondent to depolarizing steps (putative astrocytes), were unstable, or did not exhibit spontaneous activity; therefore, these recordings were discarded from the analysis.

Spontaneous excitatory postsynaptic potentials (sEPSPs) were recorded in current-clamp gap-free mode (clamped at −45 mV). Current-clamp recordings (at −60 mV) of evoked action potentials were performed by applying a repetitive current pulse (800 ms) with an incremental amplitude (20 pA).

Data analysis was performed using Clampfit 10.6 software (Molecular Devices). To visualize the distribution of the firing frequency, the number of spontaneous spikes per second in current-clamp mode was counted over a one-minute period using the threshold tool of Clampfit software and classified in bins with a width equal to 1 (corresponding to 1 Hz). The data were visualized using the frequency distribution mode of GraphPad Prism V9 software.

### HPLC analysis of dopamine content

At 80 DIV, 1-2 organoids per sample were collected and homogenized in 100 µl of 0.2 M HClO_4_. Then, the samples were centrifuged, and the supernatant was collected and spun through a 0.2 µm spin filter (Costar Spin-X, Merck) at 14000 × g at 4 °C for 1 min and loaded into an HPLC system (Thermo Scientific Ultimate 3000). The mobile phase was 12.5% acetonitrile buffer (pH 3.0, 86 mM sodium dihydrogen phosphate, 0.01% triethylamine, 2.08 mM 1-octanesulfonic acid sodium salt, and 0.02 mM EDTA). The flow rate of the mobile phase was adjusted to 1.5 ml/min. The dopamine level was calculated using a standard curve generated using external DA standards (the standard curve coefficient of determination was 0.99946). Dopamine content was then normalized to the protein concentration and is expressed in nmol/g.

### Preparation of cells for *in vivo* transplantation

For each batch of single cells, ten neurospheres at 16 DIV from each cell line (H9 and 4X) were collected and washed twice with PBS. Then, 500 µl of Accutase (supplemented with 100 µg/ml DNase) was added, and the cells were incubated for 10 min at 37 °C. The neurospheres were first pipetted with a 1 ml pipette followed by a 200 µl pipette to yield a single-cell solution. Five hundred microliters of washing medium (DMEM/F12 supplemented with 1% human serum albumin) was added, and the cells were spun down at 400×g for 5 min at room temperature. The cell pellets were resuspended at a concentration of 100,000 cells/µl in HBSS (supplemented with 100 µg/ml DNase) and kept on ice. The cell suspension was kept on ice for a maximum of three hours, after which a new batch of cells was prepared.

### *In vivo* transplantation

Adult (9 weeks old) male (225-300 g) (n= 30) NIH (NTac:NIH-*Foxn1*^*rnu*^) nude rats purchased from Taconic Biosciences A/S were grouped-housed in ventilated cages in a clean room under a 12-hr light/dark cycle with *ad libitum* access to sterile food and water. In addition to a standard rat diet, they were given peanuts to increase caloric intake. All animal experiments were conducted in accordance with the guidelines of the European Union Directive (2010/63/EU) and approved by the Danish Animal Inspectorate.

The rats were anesthetized with isoflurane (5% for induction, 2-3% for maintenance), 1.2 L/min of O_2_, and 0.6 L/min of atmospheric air, and placed in a stereotaxic frame (Stoelting) and unilaterally injected with 6-OHDA (Sigma-Aldrich A/S) (2 µl of 7 µg/µl free base in saline containing 0.02% ascorbic acid) ^49^ into the right MFB (anteroposterior (AP), -4.4; mediolateral (ML) -1.1; dorsoventral (DV), -7.6; tooth bar, 3.3) using a Hamilton syringe with a glass cannula attached. Following injection, the cannula was left in place for 5 min before being slowly retracted. The incision was sutured, and the animals were injected with buprenorphine (0.36 mg/kg) as an analgesic. Once the animals were fully awake, they were placed back into their cages with wet food and 0.009 mg/ml Temgesic in water.

Lesioning efficiency was assessed 3 weeks postsurgery using the amphetamine-induced rotation test, and animals that exhibited >5 rotations/min were used for further experiments. The selected rats were divided into 3 groups with a similar average number of amphetamine-induced rotations: the 6-OHDA lesion (no transplantation) group (n = 8), the H9 cell-transplanted group (n = 9) and the 4X cell-transplanted group (n = 9) (see Supplementary Fig. 6a, b). Four weeks after lesioning, the animals in the H9 cell-transplanted and 4X cell-transplanted groups were stereotaxically injected into the striatum (AP, +0.5; ML, -3; DV, - 4.6/4.8) with 250,000 cells of the respective cell type in a volume of 2.5 µl using a protocol similar to the one described above. All three groups were sacrificed 22 weeks postlesioning (i.e., 18 weeks after transplantation). Two transplanted rats did not complete the study and were euthanized due to health issues: one in the 4X cell-transplanted group (week 8 posttransplantation) due to a broken tail and one in H9 cell-transplanted group due to hindlimb paralysis (week 17 posttransplantation).

### Amphetamine-induced rotation test

An amphetamine-induced rotation test was performed as described previously ^50^ one week prior to transplantation to assess the effects of the lesions and 8 and 18 weeks posttransplantation. The animals were intraperitoneally (i.p.) injected with 5 mg/kg D-amphetamine and connected to a rotameter (LE 902, PanLab, Harvard Apparatus) coupled to a LE 3806 Multicounter (PanLab, Harvard Apparatus). The number of body rotations over a period of 90 min was recorded. The data are expressed as the net number of full body turns per minute, with ipsilateral rotations having a positive value and contralateral rotations having a negative value. Animals exhibiting > 5 turns/min were considered successfully lesioned. One rat had a technical issue during one of the rotation tests and was excluded from this behavioral test.

### Cylinder test

The cylinder test was used to assess paw use asymmetry three weeks postlesioning (one week prior to transplantation) and 18 weeks posttransplantation. The animals were placed in a transparent Plexiglas cylinder (height of 30 cm, diameter of 20 cm), and two mirrors were placed behind the cylinder so that the cylinder surface could be fully visualized. Spontaneous activity was video recorded for a total of 5 min. Data analysis was performed by a researcher blinded to the groups using VCL Media Player software in slow motion as previously described ^51^. Because most of the exploratory motor activity of the animals was limited to the first 2 min and there was little movement after this timepoint, activity in the first 2 min were analyzed, and activity after this time point was analyzed only if the animal exhibited fewer than 10 movements (wall touches and rears). The following behaviors were scored to determine the extent of forelimb-use asymmetry ^51^: a) independent use of the left or right forelimb when touching the wall during a full rear or landing on the floor after a rear and b) simultaneous use of both the left and right forelimb to contact the wall of the cylinder during a full rear, for lateral movements along the wall (wall stepping) and for landing on the floor following a rear. The data are presented as the percentage of time each forelimb (left or right) or both forelimbs were used relative to all movements (wall and floor).

### Immunohistochemical analysis of brain slices

The rats were killed 23 weeks after 6-OHDA-induced lesioning by an overdose of pentobarbital (50 mg/kg i.p.). During respiratory arrest, they were perfused through the ascending aorta with ice-cold saline followed by 4% cold PFA (in 0.1 M NaPB, pH 7.4). The brains were extracted, postfixed in PFA for 2 hours and transferred to 25% sucrose solution (in 0.02 M NaPB) overnight. The brains were sectioned into 35 µm thick coronal sections on a freezing microtome (Microm HM 450, Brock and Michelsen), separated into serial coronal sections (series of 8 for the striatum and the substantia nigra), and stored at -20 °C.

Immunohistochemical staining was performed on free-floating brain sections using the following primary antibodies: mouse anti-rat TH (1:4000, MAB318, Merck Millipore), rabbit anti-Girk2 (1:500, APC-006, Alomone), rabbit anti-TH (1:1000, PelFreeze), mouse IgG1 anti-CALB1 (1:5000, 28k, SWANT), mouse IgG1 anti-HNA (1:200, 151181, Abcam), goat anti-FOXA2 (1:200, AF2400), sheep anti-hCOL1A1 (1:200, R&D Systems), rabbit anti-hCOL3A1 (1:1000), rabbit anti-EN1 (1:50), and rabbit anti-LMX1A (1:5000).

Immunohistochemistry was performed as previously described ^49^ with avidin-biotin-peroxidase complex (ABS Elite, Vector Laboratories) and 3,3-diaminobenzidine (DAB) as a chromogen to visualize the signal. The sections were mounted on chrome-alum gelatin-coated slides, dehydrated, and coverslipped. The slides were analyzed using a Olympus VS120 Slide Scanner (upright widefield fluorescence) with a 20x objective.

For immunofluorescence, free-floating sections were blocked in 5% normal donkey serum in 0.25% Triton X-100 in KBPS and then incubated overnight with the selected primary antibody in 2.5% donkey serum and 0.25% Triton X-100 in KPBS at room temperature. The sections were washed with KPBS, preblocked for 10 min in 1% donkey serum and 0.25% Triton X-100 in KPBS and incubated for 2 hours with the following species-specific fluorochrome-conjugated secondary antibodies made in donkey: Alexa Fluor 488-conjugated anti-mouse IgG (1:200, Jackson ImmunoResearch), Alexa Fluor 568-conjugated anti-goat IgG (1:1000, A11057, Invitrogen), Alexa Fluor 647-conjugated anti-rabbit IgG (1:200, Jackson ImmunoResearch), Alexa Fluor 568-conjugated anti-rabbit IgG (1:1000, A10042, Invitrogen), Alexa Fluor 647-conjugated anti-mouse IgG (1:1000, A-31571, Invitrogen) and Alexa Fluor 568-conjugated anti-mouse IgG1 (1:1000, A10037, Invitrogen), Alexa Fluor 568-conjugated anti-sheep IgG (1:1000, Invitrogen). DAPI (1:2000, Sigma-Aldrich A/S) was used for nuclear staining. The sections were mounted on chrome-alum gelatin-coated slides with Dako fluorescent mounting medium.

### Microscopic analysis (TH-positive cell number and yield and graft volume)

Coronal sections (1:8) from each animal were immunostained for TH, and DA neurons in the graft were analyzed. An Olympus VS120 Slide Scanner (upright widefield fluorescence) (Bioimaging Core Facility, Aarhus University) was used to acquire images of the slides using a 20x objective. All sections with visible grafts were selected: 3-5 sections per H9 cell-transplanted animal and 4-8 sections per 4X cell-transplanted animal. The area in which the number of TH-positive cells was quantified included the striatum and globus pallidus, but TH-positive cells in the cortex and corpus callosum were not included. The images were analyzed by identifying cells in the region of interest (ROI) using QuPath software ^52^. The settings adapted for each section depending on the staining, and the following settings were used: detection image = optical density sum, requested pixel size = 0.5 µm, background radius = 15-30 µm, threshold = 0.15-0.3, median filter radius = 0-3 µm, sigma = 0.7-2 µm, minimum area = 85-130 µm^2^, maximum area = 500-1200 µm^2^, max background intensity = 2, cell expansion = 2 µm. The cells were classified by shape, including that of the cell nucleus, and that the boundaries were smoothed.

To estimate the number of cells in a full graft, the total number of TH-positive cells per animal was determined with QuPath software and multiplied by 8, and the Abercrombie method ^53^ was used to correct for double counting of cells spanning more than one section. The Abercrombie factor of each group was calculated as the average thickness per section divided by (the averaged thickness + the average TH-positive cell size). These numbers were calculated by sampling 3 sections and 18 cells per animal from 3 different animals per group. The total number of cells in a graft was calculated as the Abercrombie factor x the total number of TH-positive cells x 8. The number of surviving cells (yield) was estimated per 100,000 transplanted cells. The volume of each graft was estimated as V = A1T1 + A2T1 +… +A_n_T1, where V is estimated volume, T1 is the sampling interval of a 1/8 series (8×35 µm), and A(n) is the area TH-positive area in the section (n) ^15^.

### Statistical analysis

All statistical analyses were performed using GraphPad Prism v 9.1.1.225. One-way ANOVA or two-way ANOVA was performed, and Sidak’s test was used for post hoc analysis when appropriate. Unpaired, two-tailed t-tests were used when comparing only the grafted groups. All data are presented as the mean ± standard error of the mean (SEM) or ± standard deviation (SD) (as indicated). P<0.05 was considered significant.

## Acknowledgments

We thank Susanne Hvolbøl Buchholdt Seldrup for assistance with culturing and maintaining the pluripotent stem cell lines. We thank Gitte Ulbjerg Toft for excellent technical help with the *in vivo* rat experiments and postmortem histological analysis of the brains. We would also like to thank Per Fuglsang Mikkelsen for assistance with HPLC. We are grateful for the assistance and use of the AU Health Bioimaging Facility, Animal Facility and FACS Core Facility. Funding: This study was supported by Lundbeckfonden grant no. DANDRITE-R248-2016-2518 and the Parkinsonforeningen. MD is a partner of BrainStem—Stem Cell Center of Excellence in Neurology, funded by Innovation Fund Denmark.

## Author contributions

Conceptualization: M.D.; methodology: M.M., M.C., N.M.J., M.M.R., M.D.; formal analysis: M.M., M.C., F.F., N.M.J., J.L., E.U., I.H.K., P.Q., M.M.R., M.D.; investigation: M.M., M.C., F.F., N.M.J., J.L., E.U., I.H.K., P.Q., M.R.R., M.D.; resources: S.N., M.R.R., M.D.; writing - original draft: M.R.R., M.D.; writing - review & editing: M.C., F.F., N.M.J., J.L., E.U., P.Q., M.M.R., M.D.; visualization: M.M., M.C, N.M.J., J.L., E.U., M.R.R., M.D.; supervision: M.M.R., M.D.; funding acquisition: M.M.R., M.D. All authors read and approved the final manuscript.

