## Supplementary figures for "Enhanced Production of Mesencephalic Dopaminergic Neurons from Lineage-Restricted Human Undifferentiated Stem Cells"

Summary of content:

Supplementary Figures

Supplementary Figure 1, generation of GBX2 knockout

Supplementary Figure 2, generation of 4X knockout

Supplementary Figure 3, additional data for Fig. 2

Supplementary Figure 4, additional data for Fig. 3

Supplementary Figure 5, additional data for Fig. 4

Supplementary Figure 6, additional data for Fig. 5

Supplementary Tables

Supplementary Table 1, guide strand target sequence including PAM

Supplementary Table 2, QPCR primers

Supplementary Table 3, Nanostring probes

**Supplementary Table1:**

Guide RNA sequences used to generate knockout cell lines

| Cell line | Genes | CRISPR target sequence + PAM |
|---|---|---|
| **GBX2-/-** | GBX2 | gRNA1: GGCAGCACTACCGGCCGGTA GGG<br>gRNA2: ATGATGATGCAGCGCCCGCT GGG<br>gRNA3: GCATCATCATCAGCGACGGC GGG<br>gRNA4: AAAGGTGGAAGACGACCCGA AGG |
| **4X (GBX2-/- CDX1/2/4-/-)** | GBX2<br>CDX1<br>CDX2<br>CDX4 | gRNA4: AAAGGTGGAAGACGACCCGA AGG<br>gRNA1: CTACACCGACCACCAACGCC TGG<br>gRNA1: GTACACGGACCACCAGCGGC TGG<br>gRNA1: CCTGGGCCTTTCCGAGAGAC AGG |

**Supplementary Table 2:**

Taqman QPCR primers.

| Genes | Assay ID |
|---|---|
| OTX2 | Hs00222238_m1 |
| LMX1A | Hs00898455_m1 |
| CDX2 | Hs01078080_m1 |
| MAFB | Hs00534343_s1 |
| HOXA2 | Hs00534579_m1 |
| HOXA3 | Hs00601076_m1 |
| HOXB1 | Hs00157973_m1 |
| HOXB2 | Hs01911167_s1 |
| CNPY1 | Hs01073160_m1 |
| BARHL1 | Hs01063929_m1 |
| IRX3 | Hs01124217_g1 |
| RPL32 | Hs00851655_g1 |

**Supplementary Table 3:**

Nanostring probes.

| Genes | Probe sequence |
|---|---|
| OTX2 | GGCTGGACATTCCAGTTTTAGCCAGGCATTGGTTAAAAGAGTTAGATGGGATGATGCTCAGACTCATCTGATCAAAGTTCCGAGAGGCATAGAAGGAAAA |
| EN1 | GCAGCATTTTTGAAAAGGGAGAAAGACTCGGACAGGTGCTATCGAAAAATAAGATCCATTCTCTATTCCCAGTATAAGGGACGAAACTGCGAACTCCTTA |
| PAX8 | GACACTGTGCCCAGTGTCAGCTCCATTAATAGAATCATCCGGACCAAAGTGCAGCAACCATTCAACCTCCCTATGGACAGCTGCGTGGCCACCAAGTCCC |
| HOXA2 | CCCAAAGTTTCCCAGTCTCGCCTTTAACCAGCAATGAGAAAAATCTGAAACATTTTCAGCACCAGTCACCCACTGTTCCCAACTGCTTGTCAACAATGGG |
| HOXA4 | TGCACTTCACAAATTAATGACCATGAGCTCGTTTTTGATAAACTCCAACTACATCGAGCCCAAGTTCCCTCCCTTCGAGGAGTACGCGCAGCACAGCGGC |
| HOXB8 | GTGGTAGTATCTCGTAATAGCTTCTGTGTGTGAGCTACCGTGGATCTCCTTCCCTTCTCTTGGGGGCCGGGGGGAAAGAAAAGGATTTAAGCAAAGGCTC |
| HOXC10 | GGAAAGTTCGGCTAGTGTTCGTGTGTTTGTCGTAGCACCCAGAGCCTCCACCAAACCCTCTCCATGTCTTTACCTCCCAGTCGCTCTAAGAATCTGCTTG |
| PAX2 | TCCTCCTCCGGCAGGAACTGAACAGAACCACAAAAAGTCTACATTTATTTAATATGATGGTCTTTGCAAAAAGGAACAAAACAACACAAAAGCCCACCAG |
| PAX5 | CTCCAAGAGGAGCACACTTTGGGGAGATGTCCTGGTTTCCTGCCTCCATTTCTCTGGGACCGATGCAGTATCAGCAGCTCTTTTCCAGATCAAAGAACTC |
| FGF8 | AGAGCAACGGCAAAGGCAAGGACTGCGTCTTCACGGAGATTGTGCTGGAGAACAACTACACAGCGCTGCAGAATGCCAAGTACGAGGGCTGGTACATGGC |
| IRX3 | AGTCGCTTCTGTGGCACCCCGCATTCGCTGTGAGGTTTGTTTGTCCGGTTGATTTTGGGGGGTGGAGTTTCAGTGAGAATAAACGTGTCTGCCTTTGTGT |
| HOXA1 | CAGATAATTCTGGACCAGAGACTTGGTGCGGGGTTAACACCTTCATCCAGATTGGGTGCCAGCATACATTTTCTGGTGGGCCTTAACATCCCTCCTGCTT |
| HOXC6 | ACGTCGCCCTCAATTCCACCGCCTATGATCCAGTGAGGCATTTCTCGACCTATGGAGCGGCCGTTGCCCAGAACCGGATCTACTCGACTCCCTTTTATTC |

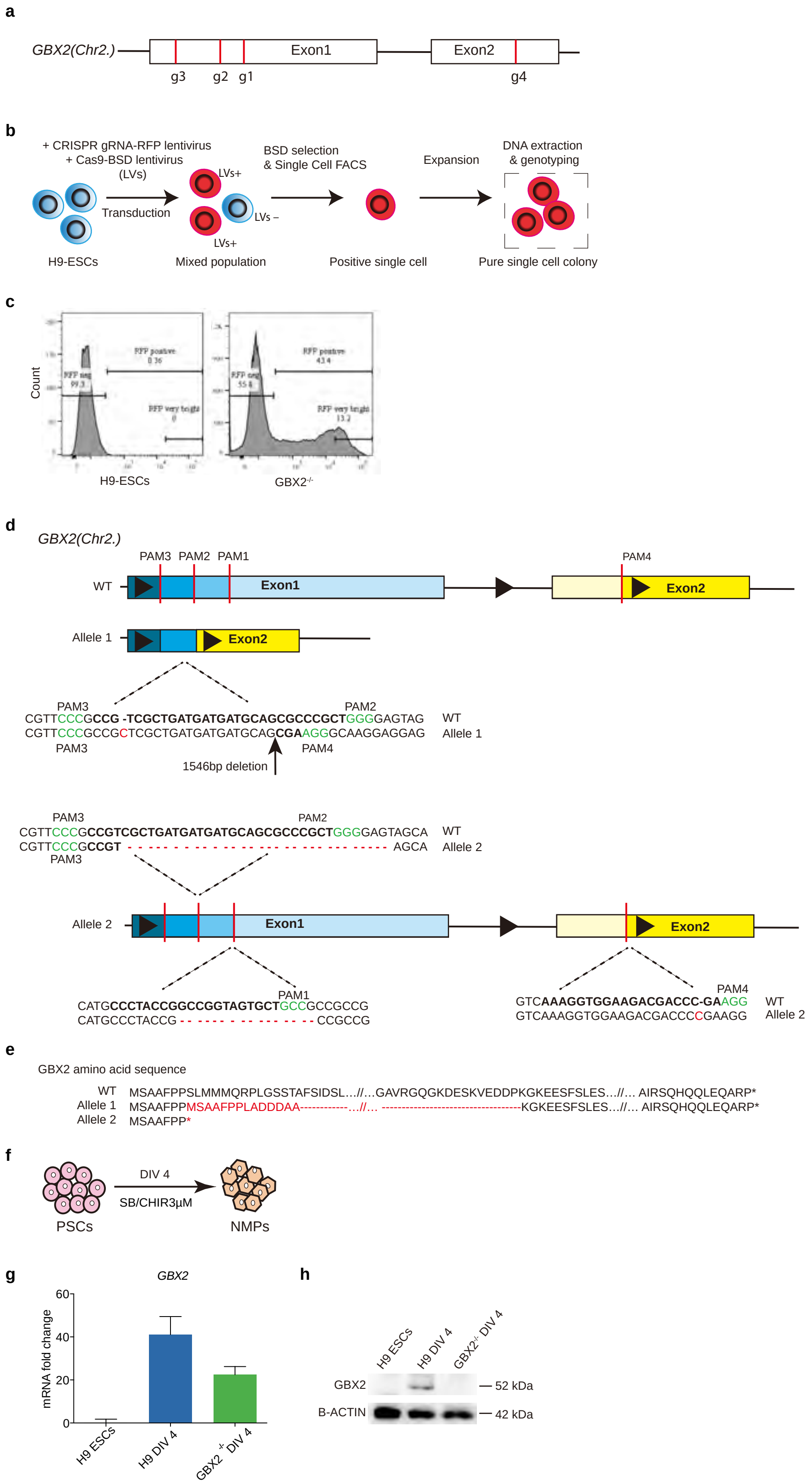


**Supplementary Figure 1: Generation of GBX2$^{-/-}$ cell line.**

**a**, Schematic diagram of the GBX2 locus, showing the guide RNA-targeted sites (g1-g4) in red lines.

**b**, Schematic overview of the CRISPR-Cas9 and single-cell-clonal selection strategy used to generate GBX2$^{-/-}$ cell line.

**c**, Single-cell FACS sorting of the transduced cells based on the RFP signal intensity. H9 hESC line is used as a negative gating control for RFP.

**d**, Mutational analysis by whole exome sequencing revealed biallelic indels mutation in the targeted GBX2 loci.

**e**, Predicted protein sequence of the mutant alleles. The out-of-frame sequences are shown in red.

**f**, Schematic diagram of the differentiation protocol used to validate the loss of GBX2 in GBX2$^{-/-}$ cell line.

**g**, RNA expression analysis of *GBX2* in H9 ESCs, H9 and GBX2$^{-/-}$ cells at DIV4.

**h**, Western blot analysis of GBX2 in H9 ESCs and neural mesoderm progenitors (NMPs) at DIV4, showing no detectable GBX2 in GBX2$^{-/-}$ cell line. β-actin shown as loading control. H9 ESCs are used as a negative control in **(g)** and **(h)**. Blasticidin (BSD).

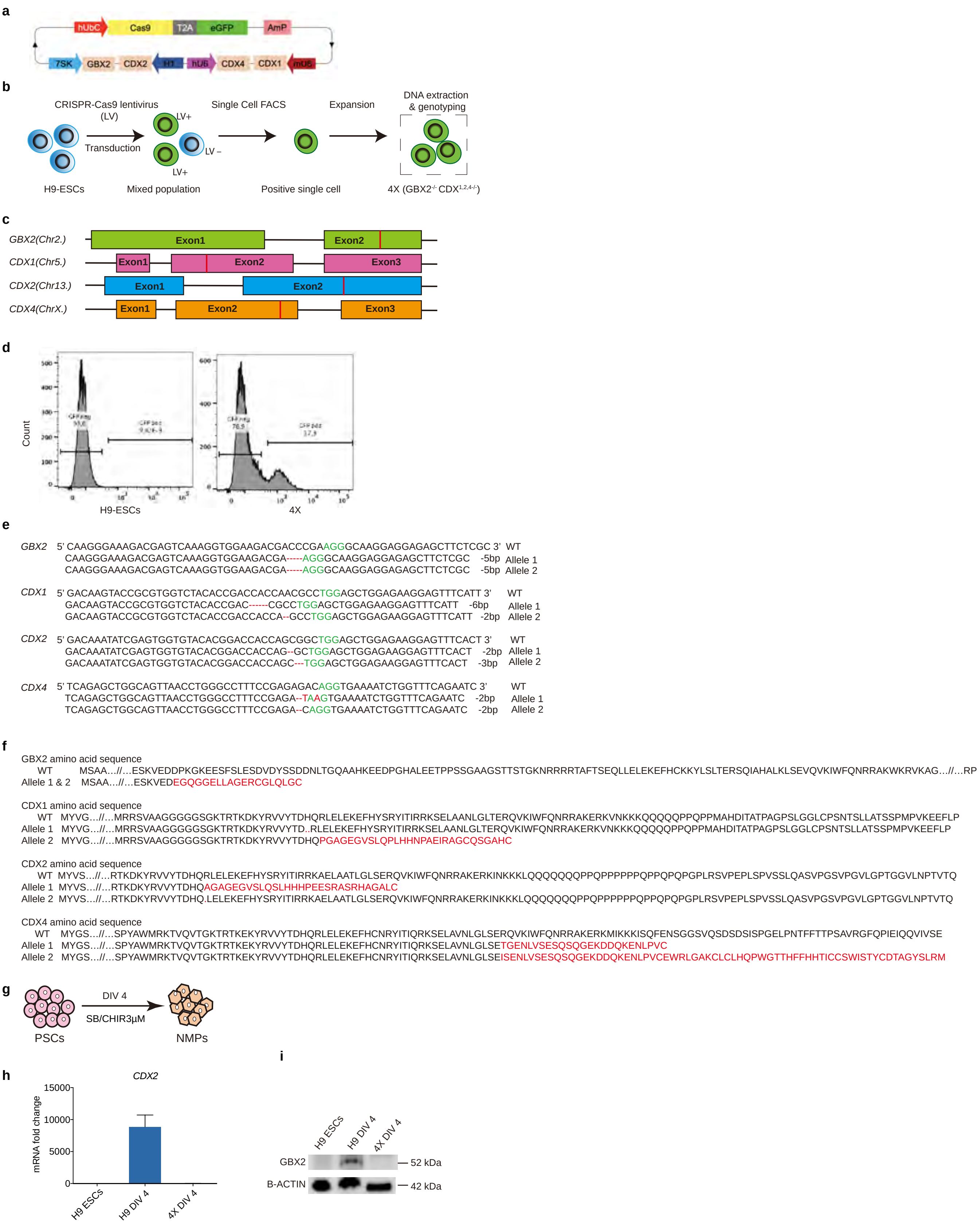


**Supplementary Figure 2: Generation of 4X cell line.**

**a**, Schema of the Cas9 lentiviral construct used to target *GBX2, CDX1, CDX2* and *CDX4*.

**b**, Schematic overview of the single cell clonal selection of CRISPR-modified hESCs.

**c**, Schematic diagram of the Cas9-targeted genes. The guide RNA-targeted sites (g1-g4) are indicated by red lines.

**d**, Single-cell FACS sorting of the transduced cells based on the GFP signal intensity. H9 hESC line is used as a negative gating control for GFP.

**e**, Mutational analysis by whole exome sequencing identified biallelic mutations in the targeted genes.

**f**, Predicted protein sequence of the mutant alleles. The out-of-frame sequences are shown in red.

**g**, Schematic diagram of the differentiation protocol used to validate the loss of CDX2 and GBX2 in 4X cell line.

**h**, RNA expression analysis of *CDX2* in H9 ESCs, H9 and 4X cells at DIV4.

**i**, Western blot analysis of GBX2 in H9 ESCs and neural mesoderm progenitors (NMPs) at DIV4, showing no detectable GBX2 protein in 4X cell line. β-actin shown as loading control. H9 ESCs are used as a negative control in **(h)** and **(i)**.

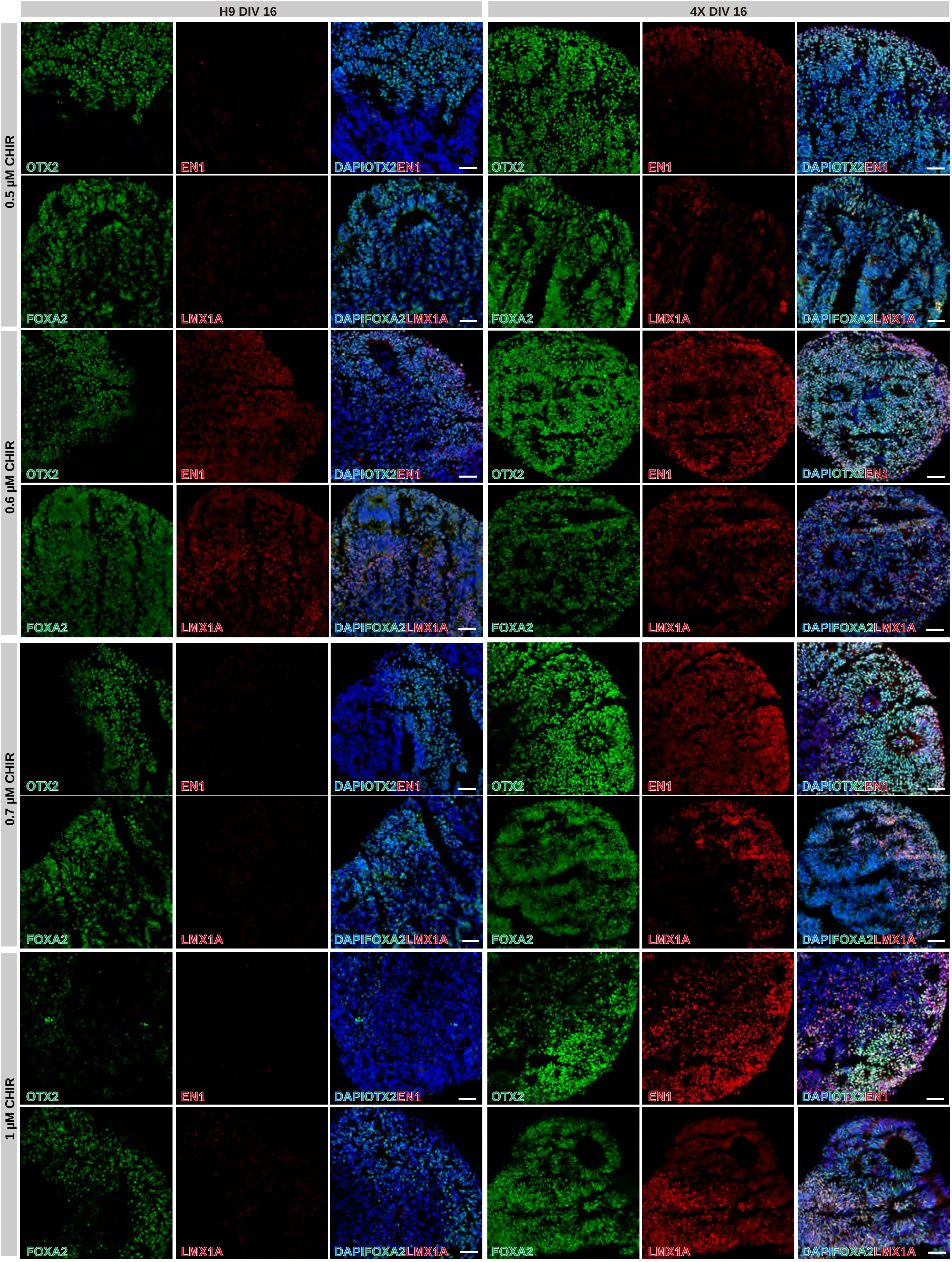


**Supplementary Figure 3: Phenotype of H9 and 4X cultures on 16 DIV with varying concentrations of GSK3i.** Representative immunofluorescence analysis identifies co-staining of OTX2/EN1 and FOXA2/LMX1A in H9 and 4X DIV16 cultures, at varying concentrations of GSK3i (0.5 µM, 0.6 µM, 0.7 µM, and 1 µM). Scale bar, 50 µm.

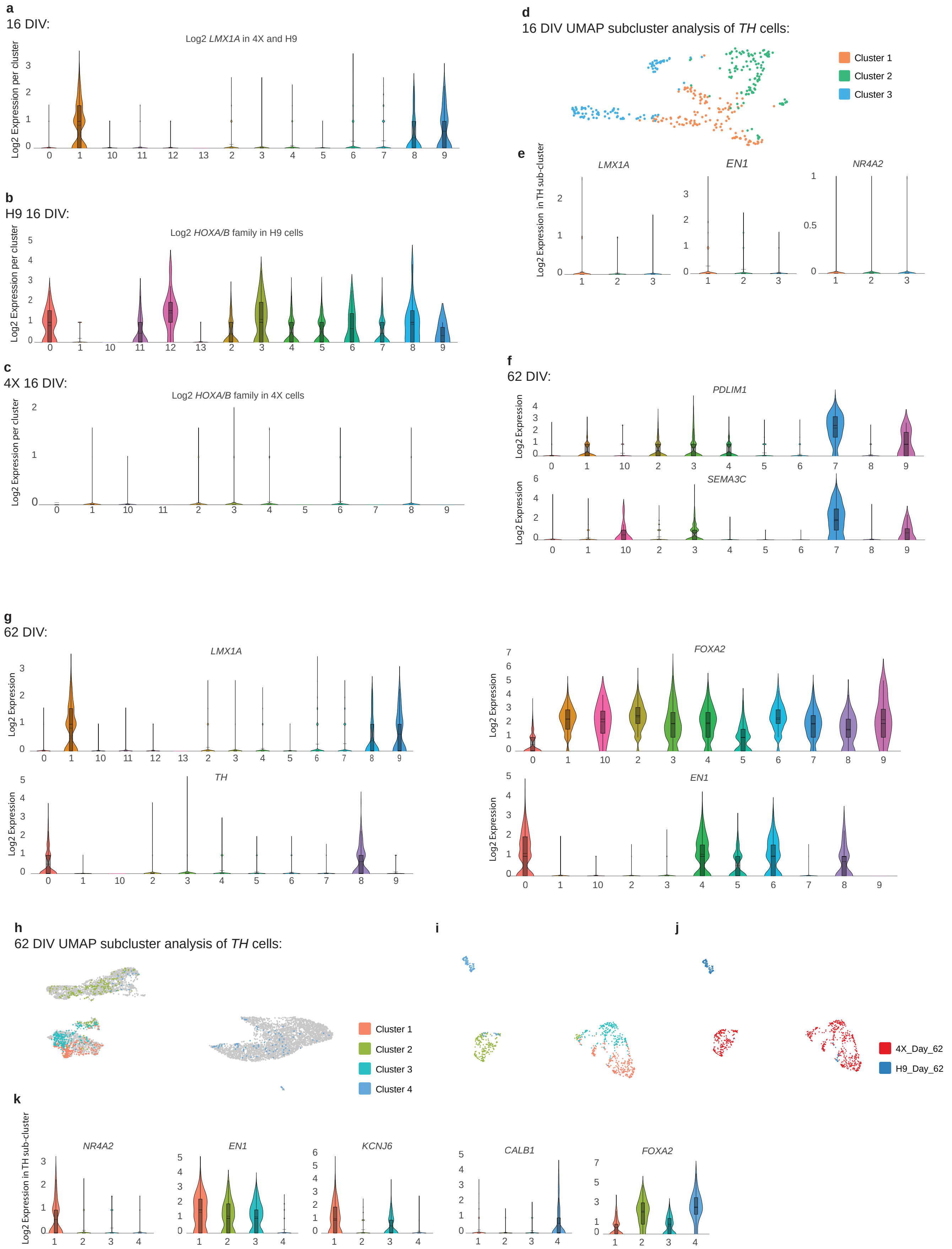


**Supplementary Figure 4: Single cell analysis on 16 DIV and 62 DIV .**

**a**, Violin plot of *LMX1A* in 4X and H9 cells across the clusters at 16 DIV.

**b**, Violin plot of *HOXA/B* family in H9 cells across the clusters.

**c**, Violin plot of *HOXA/B* family in 4X cells across the clusters.

**d**, Sub-cluster analysis of *TH* cells at 16 DIV, which results in three sub-clusters.

**e**, Violin plots of *LMX1A, EN1* and *NR4A2* in 16 DIV TH sub-clusters.

**f**, Violin plot of *PDLIM1* and *SEMA3C* in 62 DIV clusters.

**g**, Violin plot of *LMX1A, TH, FOXA2* and *EN1* in 62 DIV clusters.

**h**, Sub-cluster analysis of *TH* cells at 62 DIV, which results in 4 sub-clusters projected back onto UMAP plot.

**i**, UMAP of *TH* sub-clusters, **j**, 4X and H9 distribution within *TH* sub-clusters at 62 DIV.

**k**, Violin plot of NR4A2*, EN1, KCNJ6, CALB1* and *FOXA2* in *TH* sub-clusters at 62 DIV.

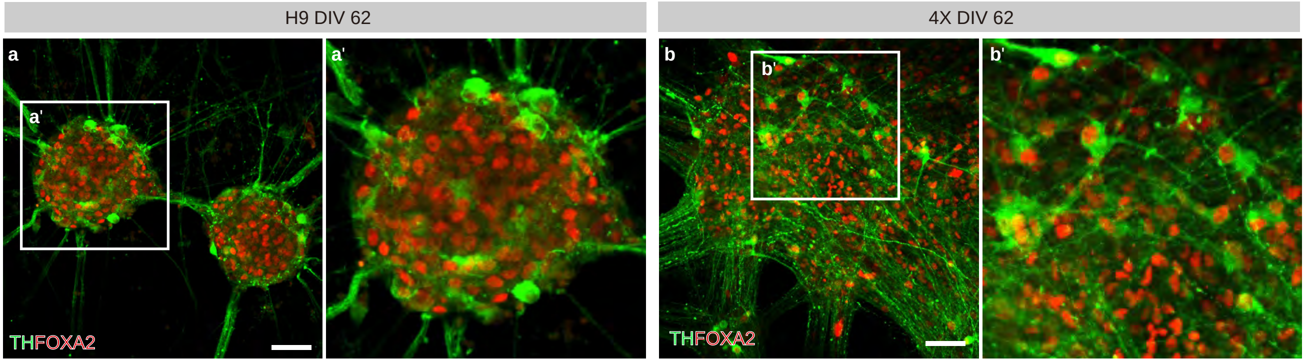


**Supplementary Figure 5: TH/FOXA2 dopaminergic neurons in H9 and 4X 2D cultures on DIV62 at 1µM concentration of GSK3i.**

**a-b**, Representative immunofluorescence analysis, revealing low and high content of double positive TH/FOXA2 cells in H9 **(a)** and 4X **(b)** DIV62 cultures, respectively. The squares in **(a-b)** outline the areas of magnification shown on **(a'-b')**. Scale bar, 50µm.

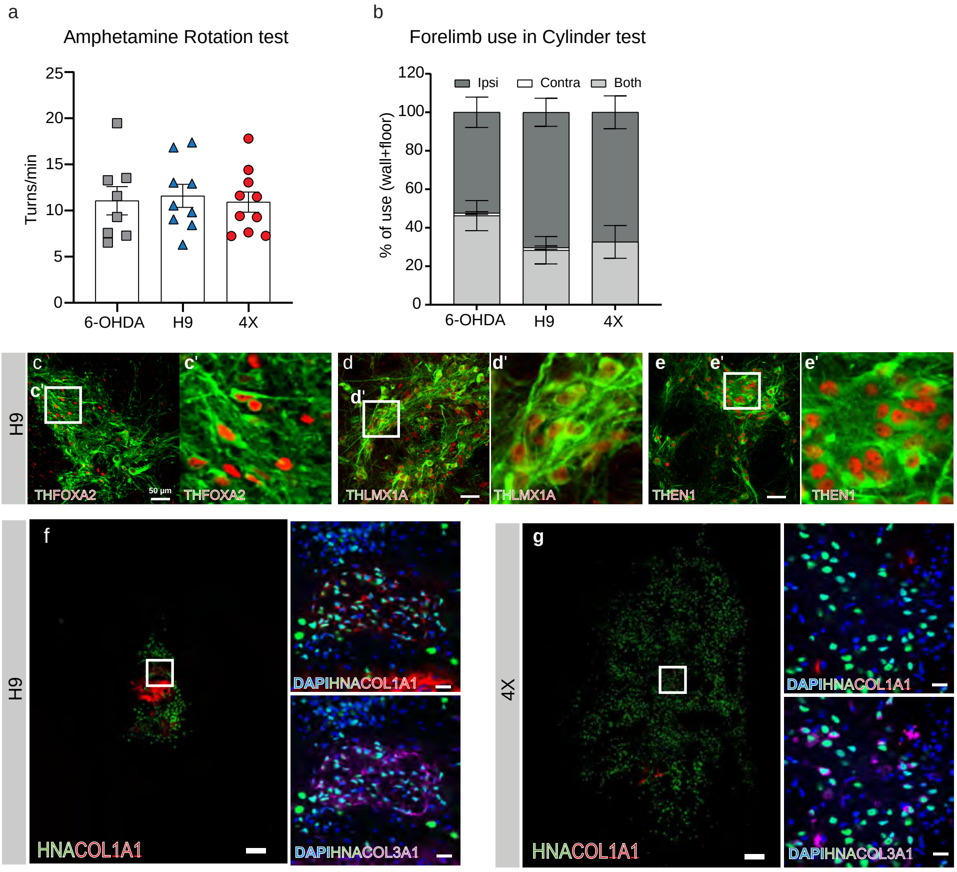


**Supplementary Figure 6: Behavior analysis of 6-OHDA-lesioned rats 3-weeks post lesion and assessment of mesDA neuron and/or vascular cell content in H9 and 4X grafts, 18 weeks post-transplantation.**

**a-b**, Amphetamine-induced rotation **(a)** and cylinder tests **(b)** of 6-OHDA-lesioned rats 3-weeks post lesion, showing comparable behavior among 6-OHDA, H9 and 4X groups, subdivided prior to transplantation. For **(a, b)**, N=8 for 6-OHDA, N=9 for H9, N=10 for 4X.

**c-e**, Representative immunofluorescence analysis identifies co-staining of TH/FOXA2, TH/LMX1A and TH/EN1 in H9 grafts, 18-weeks post-transplantation. The squares in **(c-e)** outline the areas of magnification shown on **(c'-e')**. Scale bar, 50µm.

**f-g**, Representative photomicrographs and immunofluorescence images of H9 **(f)** and 4X **(g)** grafts, co-stained with HNA/COL1A1 (left and right top panel) and HNA/COL3A1 (right bottom panel). H9 grafts have many HNA cells positive for COL1A1 and COL3A1, whereas 4X grafts exhibit no or few HNA cells positive for COL1A1 and COL3A1, respectively. DAPI was used as nuclear stain. Scale bars, 200µm (left panel) and 20µm (right top and bottom panels).
